# H2A.Z chaperones converge on histone H4 acetylation for melanoma cell proliferation

**DOI:** 10.1101/2023.11.26.568747

**Authors:** Sina Jostes, Chiara Vardabasso, Joanna Dong, Saul Carcamo, Rajendra Singh, Robert Phelps, Austin Meadows, Dan Hasson, Emily Bernstein

**Author notes:** Corresponding Author: Emily Bernstein. These authors contributed equally to this work (Chiara Vardabasso, Joanna Dong).

## Abstract

High levels of H2A.Z promote melanoma cell proliferation and correlate with poor prognosis. However, the role of the two distinct H2A.Z histone chaperone complexes, SRCAP and P400-TIP60, in melanoma remains unclear. Here, we show that individual depletion of *SRCAP*, *P400*, and *VPS72* (YL1) not only results in loss of H2A.Z deposition into chromatin, but also a striking reduction of H4 acetylation in melanoma cells. This loss of H4 acetylation is found at the promoters of cell cycle genes directly bound by H2A.Z and its chaperones, suggesting a highly coordinated regulation between H2A.Z deposition and H4 acetylation to promote their expression. Knockdown of each of the three subunits downregulates E2F1 and its targets, resulting in a cell cycle arrest akin to H2A.Z depletion. However, unlike H2A.Z deficiency, loss of the shared H2A.Z chaperone subunit YL1 induces apoptosis. Furthermore, YL1 is overexpressed in melanoma tissues, and its upregulation is associated with poor patient outcome. Together, these findings provide a rationale for future targeting of H2A.Z chaperones as an epigenetic strategy for melanoma treatment.

## INTRODUCTION

Cutaneous melanoma is the most aggressive form of skin cancer, presenting with a high UV-induced mutational load (Sample and He 2018). Understanding the driver mutations of melanoma has led to the identification of key biological targets for melanoma therapy, such as constitutively activated BRAF (BRAF^V600E/K^) and its downstream effectors MEK and ERK (Hodis et al. 2012; Czarnecka et al. 2020). The corresponding targeted therapies such as BRAF or MEK inhibitors, and more recently, immunotherapy, have significantly improved patient outcome; however, low response rates, acquired resistance, and/or adverse events limit their success (Fedorenko et al. 2015; Griffin et al. 2017; Patel et al. 2020; Long et al. 2023). In recent years, epigenetic reprogramming has emerged as a key non-genetic driver of melanoma progression and drug resistance, and offers new opportunities to investigate targetable processes (Wang et al. 2015; Strub et al. 2018; Vardabasso et al. 2015; Filipescu et al. 2023; Zhang et al. 2021; Sah et al. 2022).

We previously reported that the evolutionary conserved H2A histone variant H2A.Z is frequently amplified in melanoma (Vardabasso et al. 2015). H2A.Z has two isoforms in vertebrates, H2A.Z.1 (*H2AFZ*) and H2A.Z.2 (*H2AFV*) (Dryhurst et al. 2009), which exert distinct, yet poorly understood functions (Giaimo et al. 2019). In melanoma, both isoforms are overexpressed and correlate with poor prognosis (Vardabasso et al. 2015). Specifically, H2A.Z.2 promotes melanoma progression by recruiting the BET (Bromodomain and Extra-Terminal domain) protein BRD2 and the transcription factor (TF) E2F1 to chromatin, facilitating expression of E2F target genes and cell proliferation (Vardabasso et al. 2015). Knockdown of H2A.Z.2 induced cell cycle arrest and sensitized melanoma cells to chemo- and targeted therapies (Vardabasso et al. 2015). However, canonical histones and their variants (i.e., H2A.Z.2) are challenging drug targets due to their high degree of homology and their flat interaction surfaces that do not provide suitable docking sites for small molecules to bind. Since the histone chaperones SRCAP (Snf2-related CBP-activator protein) and P400-TIP60 are multi-subunit complexes that deposit H2A.Z into the chromatin template, and importantly, contain various domains that can potentially be targeted, we investigated their role in melanoma.

SRCAP and P400-TIP60 are ATP-dependent complexes that catalyze the nucleosomal deposition of H2A.Z-H2B dimers in place of H2A-H2B (Latrick et al. 2016a; Ruhl et al. 2006; Gévry et al. 2007a). Both complexes are named for their scaffold proteins, SRCAP and P400, respectively. While each complex has unique subunits, SRCAP and P400-TIP60 also share key subunits such as GAS41 (*YEATS4*) and YL1 (*VPS72*). Relevant to this study, YL1 directly binds to the H2A.Z-H2B dimer through its H2A.<u>Z</u>-interacting <u>d</u>omain (ZID) and is essential for H2A.Z nucleosomal deposition (Cai et al. 2005; Ruhl et al. 2006; Latrick et al. 2016b; Liang et al. 2016). In addition to H2A.Z deposition, the P400-TIP60 complex acetylates histone H4 or H2A variants via TIP60’s lysine acetyltransferase domain, a feature lacking in the SRCAP complex (Altaf et al. 2010; Yamagata et al. 2021; Numata et al. 2020).

Here, we focus on three distinct H2A.Z chaperone subunits in melanoma cells (1) the SRCAP-specific subunit SRCAP, (2) the P400-TIP60-specific subunit P400, and (3) the shared subunit YL1. Using shRNA-mediated knockdown, we investigated the consequences of losing each individual subunit on gene expression programs, H2A.Z deposition and histone H4 acetylation (H4ac) as well as cell cycle control and viability of melanoma cells. We found that H2A.Z chaperone subunits promote cell cycle progression by activating the expression of *E2F1* and its target genes by H2A.Z deposition and H4ac at their promoters. Notably, unlike H2A.Z depletion, YL1 loss not only arrests cells in G1 but also induces apoptosis, making it a potential target for melanoma.

## RESULTS

### H2A.Z chaperones are required for H2A.Z chromatin incorporation in melanoma

In an effort to characterize the H2A.Z.1 and H2A.Z.2 interactomes in melanoma cells, we previously identified all members of the SRCAP complex, and some members of the P400-TIP60 complex as H2A.Z.1 and H2A.Z.2 binding factors by quantitative mass spectrometry (Vardabasso et al. 2015) (Supp. Fig. 1A, B). Here, we sought to validate these interactions in multiple melanoma cell lines including SK-MEL-147 and 501-MEL stably expressing H2A and H2A.Z GFP fusion proteins. In doing so, we found SRCAP, P400, and/or YL1 enriched within the pulldown of GFP-H2A.Z fusion proteins compared to that of GFP-H2A control (**Fig. 1A**). While we noticed a less pronounced enrichment of P400 and its subunits, we also found that it was less readily soluble in the MNase-based chromatin purification protocol applied here (Supp. Fig. 1C) and in our mass spectrometry studies (Vardabasso et al. 2015).

**Figure 1:**
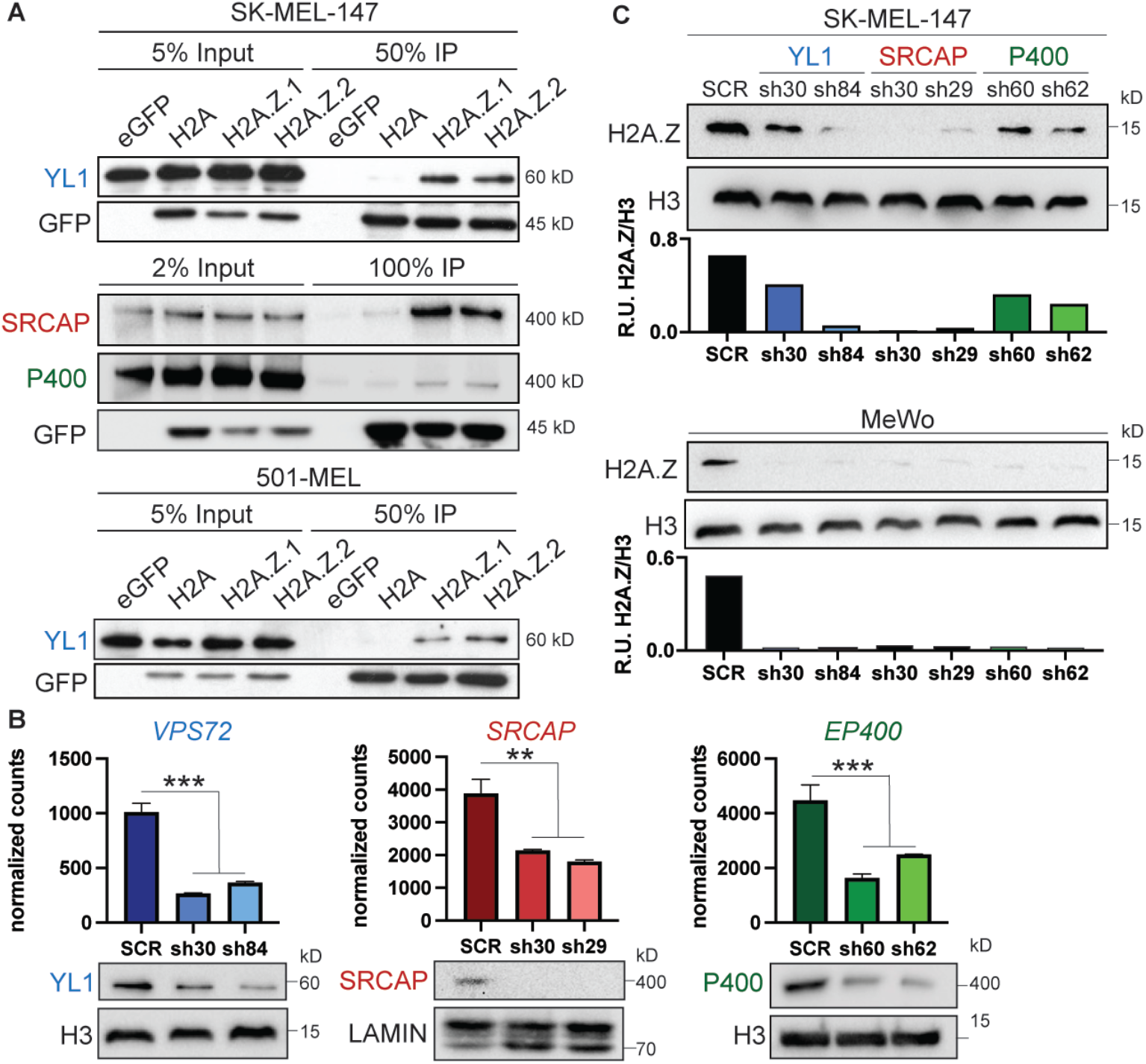
A) Anti-GFP co-IP in SK-MEL-147 and 501-MEL cells expressing GFP, GFP-H2A, GFP-H2A.Z.1 or GFP-H2A.Z.2 probed for SRCAP and P400 complex subunits. Anti-GFP blots show efficient pulldown of GFP-coupled histones. B) mRNA expression levels of *VPS72* (YL1), *SRCAP* and *EP400* (P400) as measured by RNA-seq analysis. Significance calculated using DESeq2 (*** = log2FC < -1 and padj < 0.05; ** = log2FC < -0.9 and padj < 0.05). Corresponding Western blots for YL1, SRCAP and P400 subunits shown below. H3 or LAMIN used as loading controls. C) H2A.Z Western blots of SK-MEL-147 and MeWo chromatin lysates of YL1, SRCAP and P400 knockdown samples vs. SCR control. Bar graphs show quantification of H2A.Z levels relative to H3 loading control.

We next examined H2A.Z levels in chromatin upon knockdown (KD) of YL1, SRCAP or P400 subunits. Using two independent shRNAs targeting each subunit, we were able to effectively deplete YL1, SRCAP and P400 mRNA and protein levels (**Fig. 1B**), which dramatically reduced H2A.Z levels in chromatin of SK-MEL-147 and MeWo melanoma cell lines (**Fig. 1C**). A reduction of H2A.Z following YL1 and SRCAP loss was further demonstrated in a partial CRISPR-Cas9-mediated knockout of each subunit in SK-MEL-147 cells (Supp. Fig. 1D). Thus, although primarily SRCAP subunits were enriched in our proteomic studies (Vardabasso et al. 2015); Supp. Fig. 1A, B), both SRCAP and P400-TIP60 complexes are required for H2A.Z deposition in melanoma cells.

### YL1 is overexpressed in melanoma and correlates with poor prognosis

Mining of TCGA’s cutaneous melanoma samples (363 metastatic tumor samples with mutation, CNA and expression data) (Cerami et al. 2012) revealed that *SRCAP*, *EP400* (P400) and *VPS72* (YL1) are frequently altered in melanoma at rates comparable to the defined genetic subtypes of melanoma, such as NF1 loss (**Fig. 2A**). While *SRCAP* and *EP400* are large genes with high rates of missense mutations, *VPS72* was almost exclusively altered as “mRNA high”. In line, analysis of a published microarray-based transcriptional dataset from benign nevi and primary melanomas versus human melanocytes (Talantov et al. 2005) demonstrated that *VPS72* upregulation is specific to the malignant state (**Fig. 2B**). We further performed immunohistochemical (IHC) staining of YL1 protein in benign nevi, dysplastic nevi, and primary melanomas. We observed a significant increase of YL1 in dysplastic nevi and primary melanomas (stage T1) as compared to dermal melanocytes in benign nevi (**Fig. 2C, D**). According to TCGA, the predominant alterations resulting in high *VPS72* levels in melanoma are copy number gain or amplification (**Supp.** Fig. 2A) but are not associated with any of the genetic subtypes of melanoma (**Supp.** Fig. 2B). In accordance, YL1 is highly expressed in whole cell and chromatin lysates of primary and metastatic melanoma cell lines, irrespective of their genotype, but low in normal human melanocytes (**Fig. 2E, Supp.** Fig. 2C).

**Figure 2:**
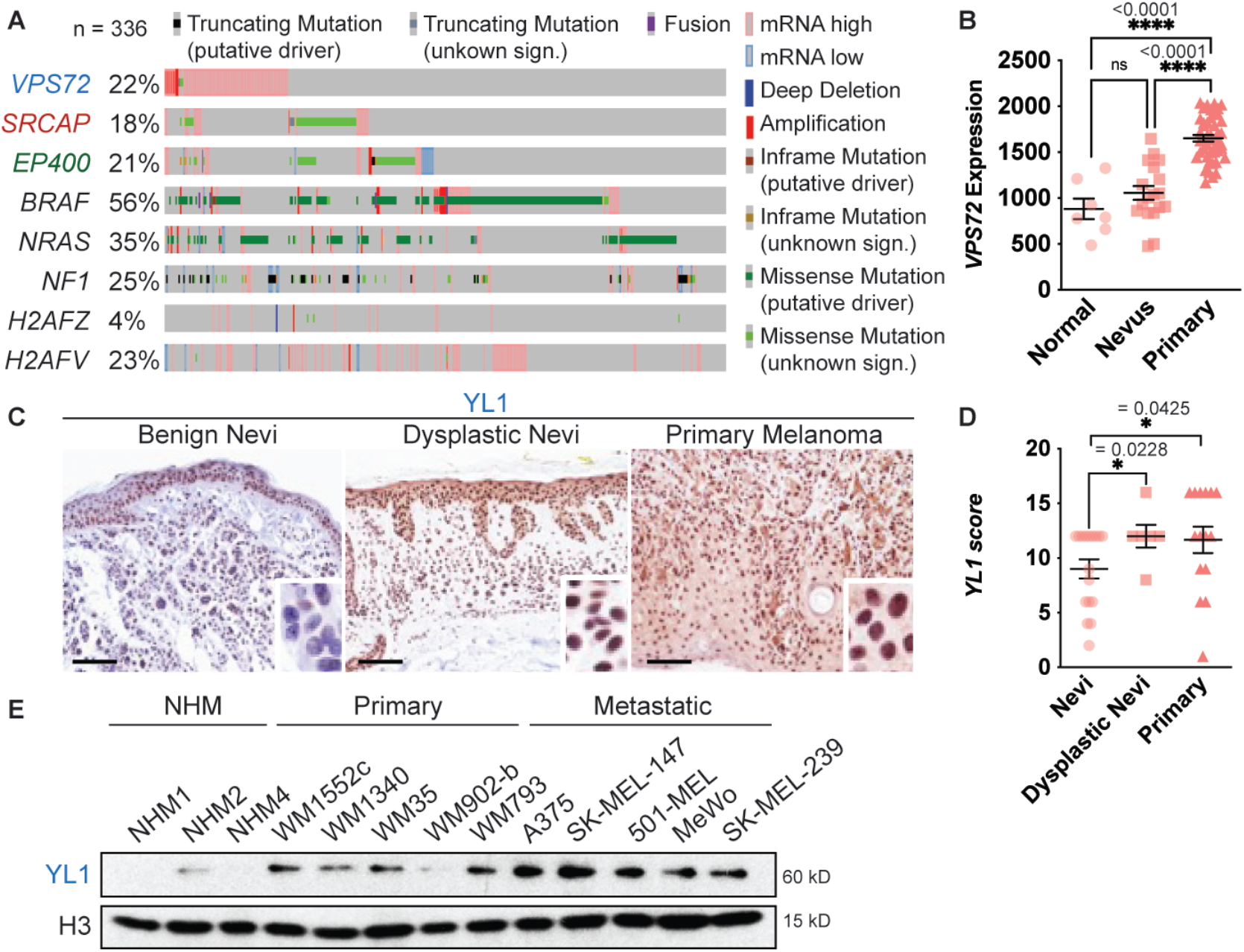
A) Alterations in Skin Cutaneous Melanoma (TCGA, PanCancer Atlas, n=363) B) *VPS72* gene expression in normal skin tissue (n=7), benign nevi (n=18) and primary melanoma (n=45) (Talantov et al. 2005). Significance was calculated using one-way ANOVA. C, D) Immunohistochemical staining of YL1 in benign nevi (n=17), dysplastic nevi (n=6) and primary melanoma samples (n=15); scoring performed by two independent pathologists. Significance was calculated using Welch’s t-test. Scale = 100 μm. Inserts are at additional 4x magnification. E) YL1 protein levels in chromatin lysate of normal human melanocytes (NHM), primary melanoma and metastatic melanoma cell lines. H3 serves as loading control.

Based on these findings, we assessed YL1 expression as a potential prognostic marker for melanoma patients. Indeed, in the TCGA cohort of primary and metastatic melanoma, high *VPS72* levels (as well as high *SRCAP* and *P400* levels) were predictive of poor survival (**Fig. 3A, Supp.** Fig. 2D). In an independent cohort of 51 primary melanoma patients (Badal et al. 2017), high *VPS72* levels were similarly predictive of poor survival (**Fig. 3A**). Here, *VPS72* expression was further able to discriminate tumors as “high risk” (*VPS72*-high) vs. ‘low risk” (*VPS72*-low) (**Fig. 3B**). The expression of *SRCAP* and *EP400* followed an opposite trend; however, their mutational status is unknown in this cohort.

**Figure 3:**
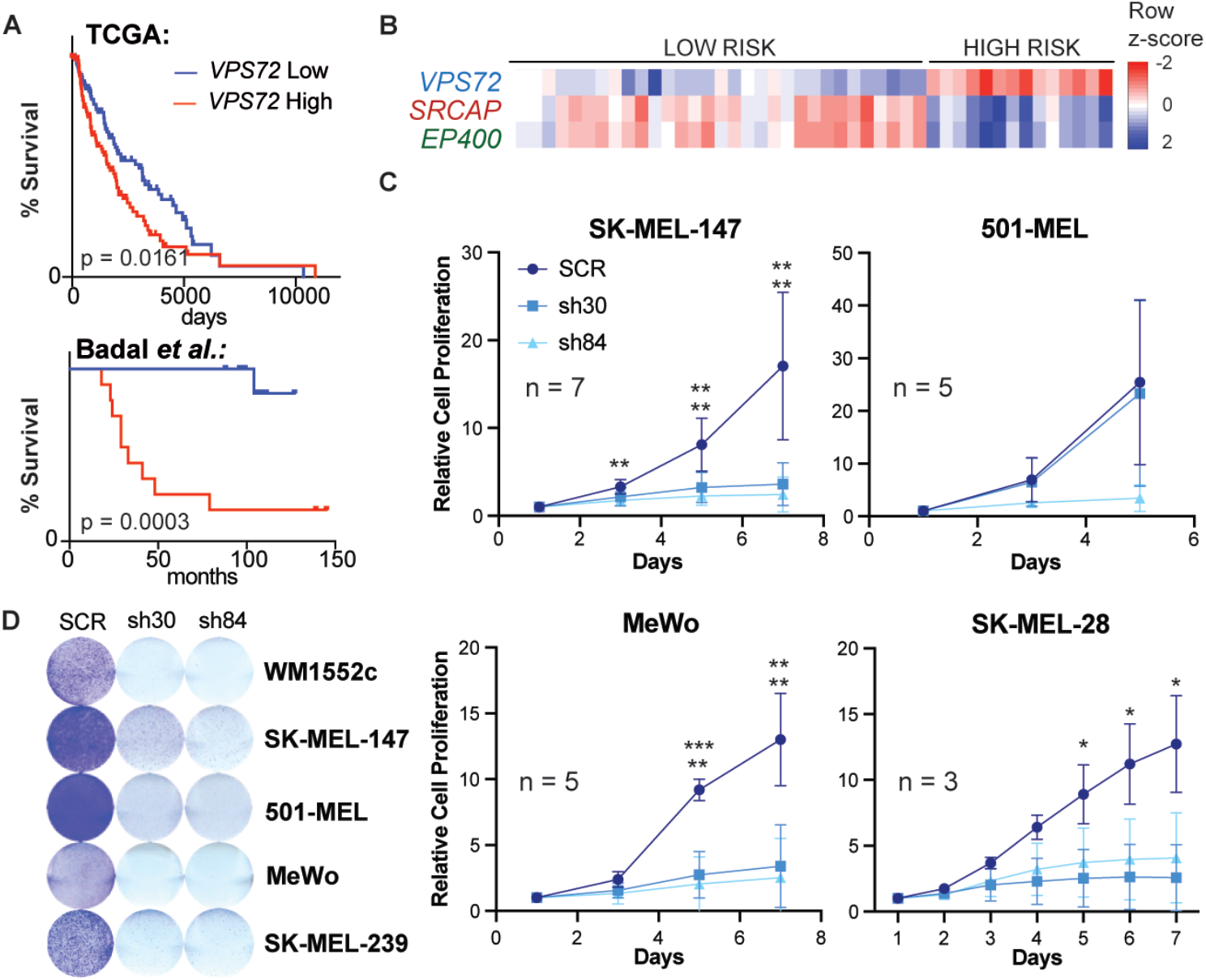
A) Survival of patients with high vs. low *VPS72* expression (divided by highest and lowest quartile) in melanoma cohorts. Upper panel = primary and metastatic melanoma (n=228, TCGA), lower panel = primary melanoma (n=44, (Badal *et al*. 2017)), significance calculated with log-rank test. B) Heatmap of expression levels of *VPS72*, *SRCAP* and *EP400* in patients with primary melanoma stratified by risk group (as defined in Badal et al. 2017). C) Proliferation of melanoma cell lines after YL1 knockdown (sh30, sh84) compared to scrambled (SCR) control over a time course of up to 7 days. Error bars indicate mean and SD. Significance calculated using 2-way ANOVA. Only significant values shown. D) Crystal violet staining of melanoma cell lines at 7 days post-knockdown with YL1 shRNAs (sh30, sh84) compared to scrambled (SCR) control.

These findings highlight that the H2A.Z chaperone subunit YL1 is overexpressed in melanoma and suggest that elevated YL1 levels may promote tumor development. To investigate this, we analyzed the effect of YL1 KD on melanoma cell proliferation *in vitro*. Indeed, we observed a significant reduction of proliferation in melanoma cell lines of distinct genetic backgrounds over the course of up to seven days (**Fig. 3C**), which was confirmed by crystal violet staining at seven days post-infection (**Fig. 3D**). We observed a comparable reduction in melanoma cell growth after SRCAP or P400 KD (**Supp.** Fig. 3A-B), suggesting that multiple H2A.Z chaperone subunits are required for melanoma cell proliferation.

### YL1, SRCAP and P400 loss results in downregulation of cell cycle-associated genes

To further assess the similarities and differences between YL1, SRCAP and P400 subunits at the transcriptomic level, we performed RNA-sequencing (RNA-seq) analysis in SK-MEL-147 cells at six days post-infection with YL1, SRCAP and P400 shRNAs. We chose this timepoint as the cells showed signs of cellular stress yet were viable enough to collect material for RNA-seq. Principal component analysis (PCA) showed that KD samples clustered separately from the controls with SRCAP KD samples showing the strongest separation (**Fig. 4A**). Interestingly, as a common subunit of both SRCAP and P400-TIP60 complexes, YL1 KD clustered between the P400 and SRCAP KD samples in the PCA.

**Figure 4:**
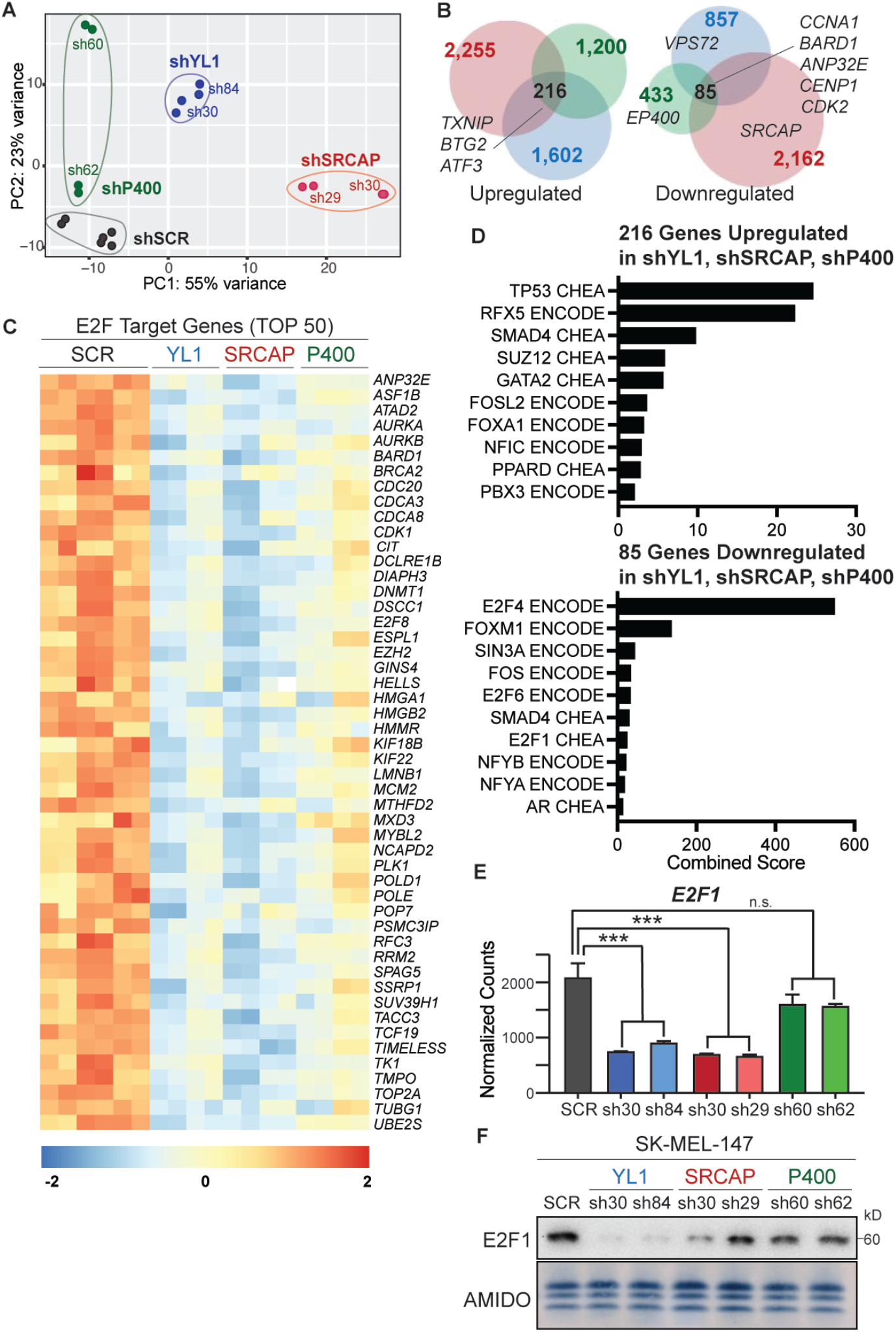
A) PCA analysis of RNA-seq samples of P400, YL1 and SRCAP knockdown samples and SCR controls. B) Venn diagrams depicting overlap of differentially expressed genes in YL1, SRCAP and P400 knockdown cells. C) Heatmap showing normalized counts of top 50 downregulated E2F target genes (as identified by GSEA) in YL1, SRCAP and P400 knockdown samples compared to SCR control. D) ChEA and ENCODE enrichment analysis of genes commonly up- or down-regulated in YL1, SRCAP and P400 knockdown samples. E) mRNA expression levels of *E2F1* in YL1, SRCAP and P400 knockdown samples compared to SCR control as measured by RNA-seq. Significance calculated using DESeq2 (*** = log2FC < -1 and padj < 0.05). F) Western blot demonstrating downregulation of E2F1 in YL1, SRCAP and P400 knockdown cells. Amido black staining of histones serves as loading control.

Next, we assessed whether KD of YL1, SRCAP or P400 would affect the gene expression of the other complex subunits (**Supp.** Fig. 4A). While none of these expression changes reached significance in our DESeq2 analysis (Ilog2FCI ≥ 0.75, padj < 0.05, **Supp. Table S1**), some partnering subunits were mildly downregulated following the KD of YL1, SRCAP or P400. Nonetheless, we did observe that the KD of YL1, SRCAP or P400 altered the protein levels of partnering subunits in chromatin, irrespective of whether they were transcriptionally downregulated or not (**Supp.** Fig. 4B). For example, YL1 KD reduced SRCAP and the SRCAP-specific subunit ZNHIT1, SRCAP KD reduced YL1, GAS41, ZNHIT1 and P400, and P400 KD reduced YL1, GAS41 and SRCAP protein levels in chromatin. This suggests that either the stability of the H2A.Z chaperone complexes depends on specific subunits (e.g. the scaffolding subunits) and/or that particular subunits are required for recruitment of the complexes to chromatin.

Given the above, as well as defects in proliferation, we hypothesized that YL1, SRCAP and P400 KD might have similar consequences on gene expression. In total, we identified 1,602 (YL1 KD), 2,255 (SRCAP KD) and 1,200 (P400 KD) upregulated and 857 (YL1 KD), 2,162 (SRCAP KD) and 433 (P400 KD) downregulated genes using an absolute |log2FC| ≥ 0.75, padj < 0.05 (**Fig. 4B**). Of those, 216 genes were commonly up- and 85 genes commonly down-regulated across YL1, SRCAP and P400 KDs (**Fig. 4B**). Despite a substantial number of deregulated genes, unchanged levels of RNA Pol II Ser5 or Ser2 phosphorylation suggest that transcription initiation or elongation processes were not globally affected by YL1, SRCAP or P400 KD (**Supp.** Fig. 5A). Gene set enrichment analysis (GSEA) showed that genes downregulated in YL1, SRCAP and P400 KD were significantly enriched for E2F Targets, G2M Checkpoint, Mitotic Spindle, and MYC Targets (**Fig. 4C, Supp.** Fig. 5B). In line, E2F was among the top enriched transcription factor signatures within the overlap of genes downregulated following YL1, SRCAP or P400 KD (**Fig. 4D**). To test which E2F family member was responsible for this signature, we compared transcript levels of E2F1-8 across all KD samples. Among the E2F members, *E2F1* was both highly expressed and downregulated in all YL1, SRCAP and P400 KD samples, with the strongest downregulation on mRNA level observed in SRCAP and YL1 KDs (**Fig. 4E**, **Supp.** Fig. 5C). This was further confirmed at the protein level, where we observed the strongest reduction of E2F1 in chromatin of YL1 KD samples (**Fig. 4E**). The observation that YL1 functions as a common subunit within both the SRCAP and P400-TIP60 complexes provides a plausible rationale for its pronounced capacity to induce transcription of E2F1.

Among significantly upregulated signatures in YL1, SRCAP and P400 KD were P53 Pathway and EMT (**Supp.** Fig. 5B, D). Induction of P21 (a primary target of P53) following H2A.Z depletion has previously been described (Gévry et al. 2007a) and was also seen in this study (**Supp.** Fig. 5D; *CDKN1A*). Upregulation of P53 protein levels in SK-MEL-147 (P53 wildtype) following YL1 KD was further demonstrated by Western blot analysis (**Supp.** Fig. 5E). Of note, in SK-MEL-28 cells, which are P53-mutant (Avery-Kiejda et al. 2011), YL1 KD led to a comparable reduction in proliferation to that of SK-MEL-147 cells (**Fig. 3C**). This suggests that P53 and its downstream effectors are not solely responsible for the observed proliferation impairment. To summarize, KD of the H2A.Z chaperone subunits YL1, SRCAP and P400 downregulates E2F1 and its target genes, resulting in reduced proliferation of melanoma cells, akin to KD of H2A.Z.2 (Vardabasso et al. 2015).

### H2A.Z chaperone subunits directly bind to *E2F1* and its targets

Next, we performed ChIP-seq analysis of YL1 and the SRCAP-specific subunit ZNHIT1 to identify common and differential genomic binding sites of the two chaperone subunits. To identify direct target genes of these chaperones via H2A.Z deposition, we further integrated our H2A.Z ChIP-seq dataset (Vardabasso et al. 2015), with histone post-translational modification (PTM) profiling and ATAC-seq (Fontanals-Cirera et al. 2017; Carcamo et al. 2022) all performed in SK-MEL-147 melanoma cells. Interestingly, clustering of YL1 and ZNHIT1 ChIP-seq data with H2A.Z ChIP-seq revealed that the majority of regions were H2A.Z-high, but YL1 and/or ZNHIT1-low (**Fig. 5A, “**Cluster 1” = 24,427 peaks, **Supp. Table S2**). These sites were almost exclusively distal intergenic regions, of which a large proportion were annotated as active (H3K4me1+, H3K27ac+) or weak/poised enhancers (H3K4me1+, H3K27ac-) (**Fig. 5B, Supp.** Fig. 6A). The weak signal for YL1, ZNHIT1 and ATAC in Cluster 1 is suggestive of a low level of histone turnover at these sites. In contrast, the majority of H2A.Z, YL1 and ZNHIT1-high regions (**“**Cluster 2” = 6,324 peaks) were mostly located at active promoters (H3K4me3+, H3K27ac+) with highly accessible chromatin, suggestive of active transcription and high turnover of H2A.Z (**Fig. 5B, Supp.** Fig. 6A). Of note, H2A.Z function depends on its PTMs; acetylated H2A.Z is associated with active transcription, while ubiquitinated H2A.Z is found predominantly at bivalent or poised enhancers (Colino-Sanguino et al. 2021). This may correlate with its role at these distinct clusters (e.g. Cluster 1 with unacetylated H2A.Z or H2A.Zub, and Cluster 2 with H2A.Zac).

**Figure 5:**
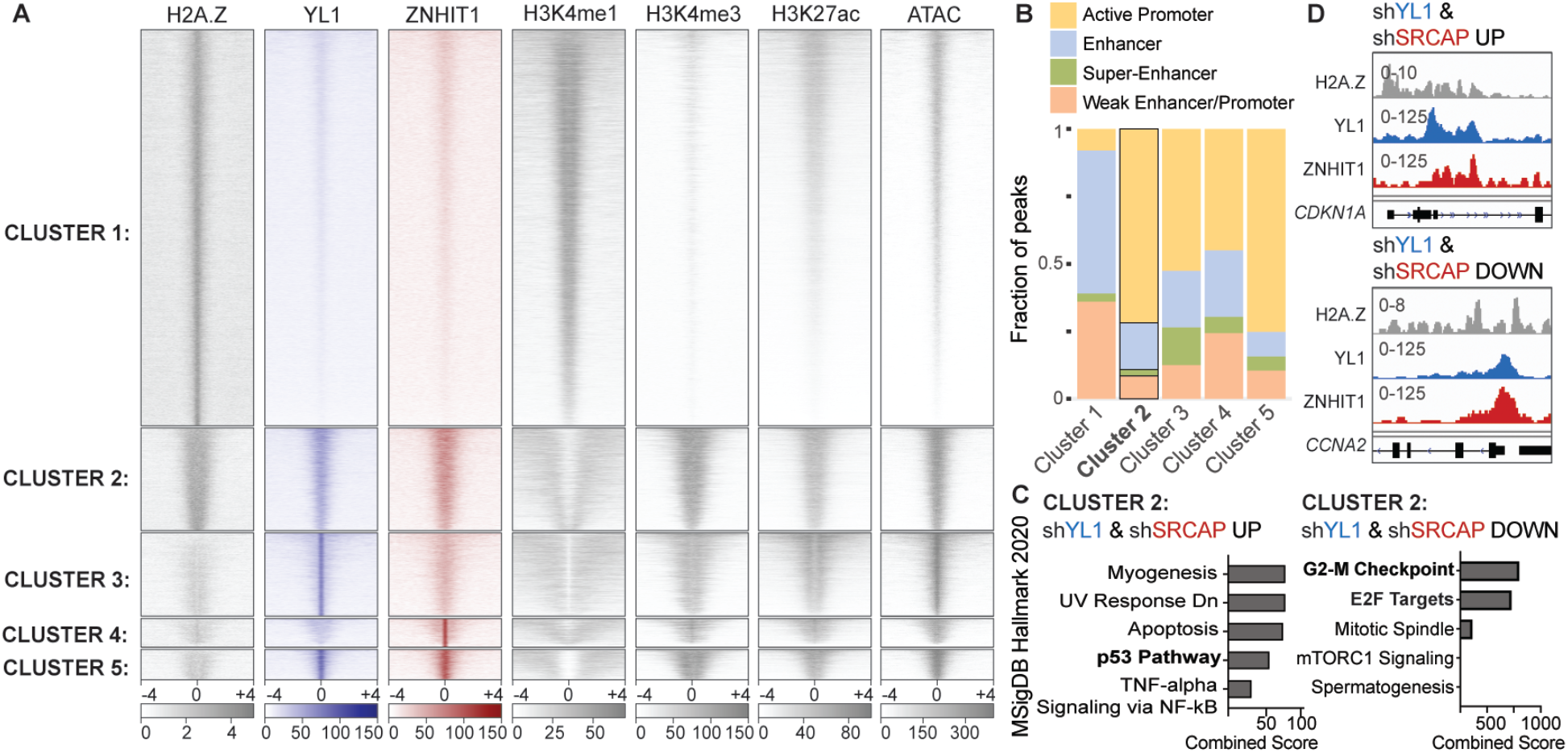
A) Heatmap of H2A.Z, YL1, ZNHIT1, H3K4me1, H3K4me3, H3K27ac and ATAC ChIP-seq signal in SK-MEL-147 cells sorted by cluster. Signal plotted around peak center. Cluster 1 = H2A.Z-High, Cluster 2 = H2A.Z + YL1 + ZNHIT1-High, Cluster 3 = YL1 High, Cluster 4 = ZNHIT1 High, Cluster 5 = YL1 + ZNHIT1 High. B) Genomic annotation of ChIP-seq peaks in Cluster 1-5. C) Enrichment analysis of genes upregulated after YL1 and SRCAP knockdown and in Cluster 2 (bound by H2A.Z, YL1 and ZNHIT1). D) Enrichment analysis of genes downregulated after YL1 and SRCAP knockdown and in Cluster 2 (bound by H2A.Z, YL1 and ZNHIT1). E) Genome Browser Tracks of H2A.Z, YL1 and ZNHIT1 ChIP-seq at *CDKN1A* (P53 target) and *CCNA2* (E2F target) gene promoters.

While we attempted to identify regions of H2A.Z deposition exclusive to the P400-TIP60 complex (i.e., not shared by SRCAP) that would be H2A.Z- and YL1-high, but ZNHIT1-low, we did not find such regions (data not shown). This suggests redundancy between SRCAP and P400-TIP60 complexes at sites of H2A.Z deposition. On the other hand, we identified regions that were H2A.Z-low but showed enrichment for YL1 (**“**Cluster 3” = 5,146 peaks), ZNHIT1 (**“**Cluster 4” = 1,765 peaks) or both (**“**Cluster 5” = 1,873 peaks) (**Fig. 5A**). Since these regions were located at active promoters and enhancers (**Fig. 5B, Supp.** Fig. 6A), it remains unclear why H2A.Z signal is low at these regions and whether YL1 and ZNHIT1 subunits may have H2A.Z-independent roles at these sites.

Notably, Gene Ontology analysis revealed that only Cluster 2, which includes all peaks bound by both H2A.Z and its chaperone subunits, is enriched for cell cycle-associated signatures (**Supp.** Fig. 6B). Other Clusters showed enrichment for (1) neuronal processes like axonogenesis (Cluster 1, H2A.Z High; Cluster 4, ZNHIT1 High), (2) actin cytoskeleton organization (Cluster 3, YL1 High) and (3) RNA processing (Cluster 5, YL1 and ZNHIT1 High). Given the role of SRCAP mutations and H2A.Z variants in neurodevelopmental disorders and neural crest development (Hood et al. 2012; Rots et al. 2021; Shi et al.; Greenberg et al. 2019), an enrichment for axonogenesis-related genes in Clusters 1 and 4 is intriguing. However, none of these gene sets were deregulated in our RNA-seq analysis, suggesting that Cluster 1, for example, may include inactive enhancers that are only active in specific cellular contexts.

Next, we overlapped genes deregulated by YL1 and SRCAP KD with the promoter peaks of Clusters 1-5 to identify direct YL1 and SRCAP target genes. Not surprisingly, we observed the highest overlap for Cluster 2, which are H2A.Z, YL1 and ZNHIT1-bound regions (**Supp.** Fig. 6C**, Supp. Table S2**). Further, Cluster 2 differentially expressed genes (DEGs) showed the most significant enrichment for P53 Pathway (upregulated in RNA-seq) and E2F targets (downregulated in RNA-seq) as identified by ChEA (ChIP enrichment analysis (Chen et al. 2013)) (**Fig. 5C, D, Supp.** Fig. 6D). Together, these findings suggest a role for H2A.Z chaperone subunits in driving expression of *E2F1* and its downstream effectors, but also in suppressing the expression of P53 target genes via H2A.Z deposition.

### H2A.Z chaperones promote transcription of E2F target genes through H2A.Z deposition and acetylation of H2A.Z and H4

We next investigated whether inhibition of H2A.Z deposition via chaperone KD altered the chromatin landscape contributing to the differential gene expression we observed. Since the P400-TIP60 complex can acetylate H2A and H4 histone tails via TIP60’s lysine acetyltransferase domain (Altaf et al. 2010a), we examined histone acetylation upon KD of H2A.Z chaperone subunits. By performing Western blot analysis following YL1, SRCAP and P400 KD, we found that loss of each individual chaperone subunit reduced levels of H2A.Z and H4 acetylation, with the strongest effects on H4ac observed for H4K16ac in both melanoma cell lines tested (**Fig. 6A**). As expected, by knocking down TIP60, we observed decreased H4 acetylation, but also a reduction of H2A.Z protein levels in chromatin lysate of SK-MEL-147 cells (**Supp.** Fig. 7A). A role for TIP60 in stimulating H2A.Z exchange has previously been described (Choi et al. 2009).

**Figure 6:**
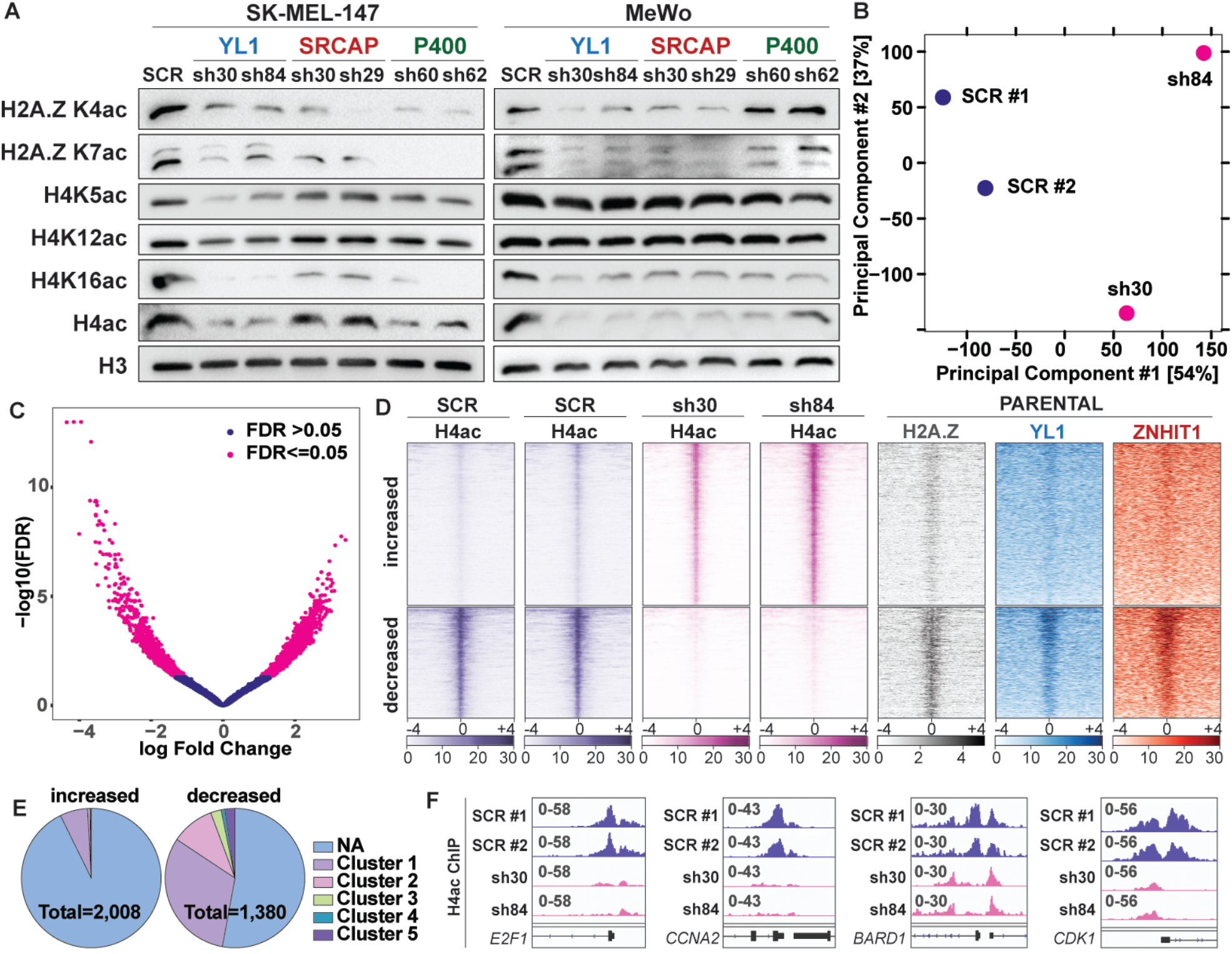
A) Western blots of H2A.Z and H4 acetylation in chromatin lysates of YL1, SRCAP and P400 knockdown samples compared to SCR control. H3 serves as loading control. B) PCA of H4ac ChIP-seq in YL1 knockdown samples (sh30, sh84) and SCR controls. C) Volcano plot displaying differential H4ac ChIP-seq peaks in YL1 knockdown samples vs. SCR controls. D) Heatmap of H4ac, H2A.Z, YL1 and ZNHIT1 ChIP-seq signal in SK-MEL-147 cells clustered by regions that gain H4ac signal (increased = 2,008) and regions that lose H4ac signal (decreased = 1,380). E) Annotation of H4ac increased and decreased regions by Cluster, see Fig. 3A for Cluster information. NA= not bound by H2A.Z or chaperone subunits. F) Genome Browser tracks of H4ac ChIP-seq at promoters of Cluster 2 genes *E2F1, CCNA2, BARD1* and *CDK1*.

To address whether the loss of H4 acetylation contributed to the downregulation of E2F targets and G2-M checkpoint genes, we next performed H4ac (Tetra-ac, H4K5ac/K8ac/K12ac/K16ac) ChIP-seq analysis in control and YL1 KD cells. PCA (principal component analysis) and correlation heatmap showed that YL1 KD samples clustered separately from SCR controls (**Fig. 6B, Supp.** Fig. 7B). In total, we identified 3,382 differential H4ac peaks, of which 2,008 were increased and 1,380 were decreased (**Fig. 6C, D, Supp. Table S3**). Next, we assessed the chromatin regions at which H4 acetylation changes occurred, by clustering them with histone modifiction profiles of promoters (H3K4me3) and enhancers (H3K4me1) and observed that H4ac decreased regions resembled mostly active promoters and enhancers, whereas H4ac increased regions were annotated as weak/poised enhancers / promoters (**Supp.** Fig. 7C, see “All regions”). Intriguingly, the majority of H4ac increased peaks were not bound by H2A.Z or H2A.Z chaperone subunits, while H4ac decreased peaks displayed enrichment of H2A.Z, YL1 and ZNHIT1 binding, suggesting that H2A.Z deposition strictly correlated with H4 acetylation (**Fig. 6D**). In fact, more than one third of the H4ac decreased peaks belonged to Cluster 1 (H2A.Z high) and 2 (H2A.Z+YL1+ZNHIT1 high) (**Fig. 6E**), mostly annotated as active promoters and enhancers (**Supp.** Fig. 7C, “Cluster 1” and “Cluster 2”). Moreover, the associated genes were enriched for G2-M and E2F targets (**Supp.** Fig. 7D). Of those, 69 genes (of which 48 genes had a peak in their promoter region) were also downregulated after YL1 KD, implying them as direct target genes. As expected, these genes included *E2F1* itself, as well as downstream effectors and cell cycle regulators like *CCNA2*, *BARD1* or *CDK1* (**Fig. 6F**). Together, these data highlight the importance of H2A.Z and its chaperones in regulating melanoma cell cycle progression by promoting a permissive, open chromatin structure at E2F target genes through H2A.Z deposition as well acetylation of histones H2A.Z and H4.

### YL1, but not H2A.Z.1 or H2A.Z.2 knockdown induces apoptosis in melanoma cells

Our data demonstrates that H2A.Z chaperones regulate E2F target and cell cycle-related genes by mediating H2A.Z deposition and H4 acetylation and that KD of H2A.Z chaperone subunits hinders melanoma cell proliferation. We aimed to further investigate these proliferation defects with a focus on YL1, which is overexpressed in melanoma samples and whose overexpression correlates with poor survival (**Fig. 2,3**). In line with cell proliferation data, the YL1 shRNAs generally induced a G1 cell cycle arrest with concomitant decreased number of cells in S phase in the melanoma cell lines analyzed including a primary melanoma (WM1552C (BRAF^V600E^)) and three metastatic melanoma lines of distinct genetic backgrounds (501-MEL (BRAF^V600E^); SK-MEL-147 (NRAS^Q61R^); MeWo (NF1^Q1136^)) (**Fig. 7A**). In addition to cell cycle arrest, we further observed a significant induction of apoptosis upon YL1 KD (**Fig. 7B**). Notably, KD of H2A.Z alone resulted in cell cycle arrest, but not apoptosis, indicating a distinction between H2A.Z and YL1 KD regarding apoptosis (Vardabasso et al. 2015).

**Figure 7:**
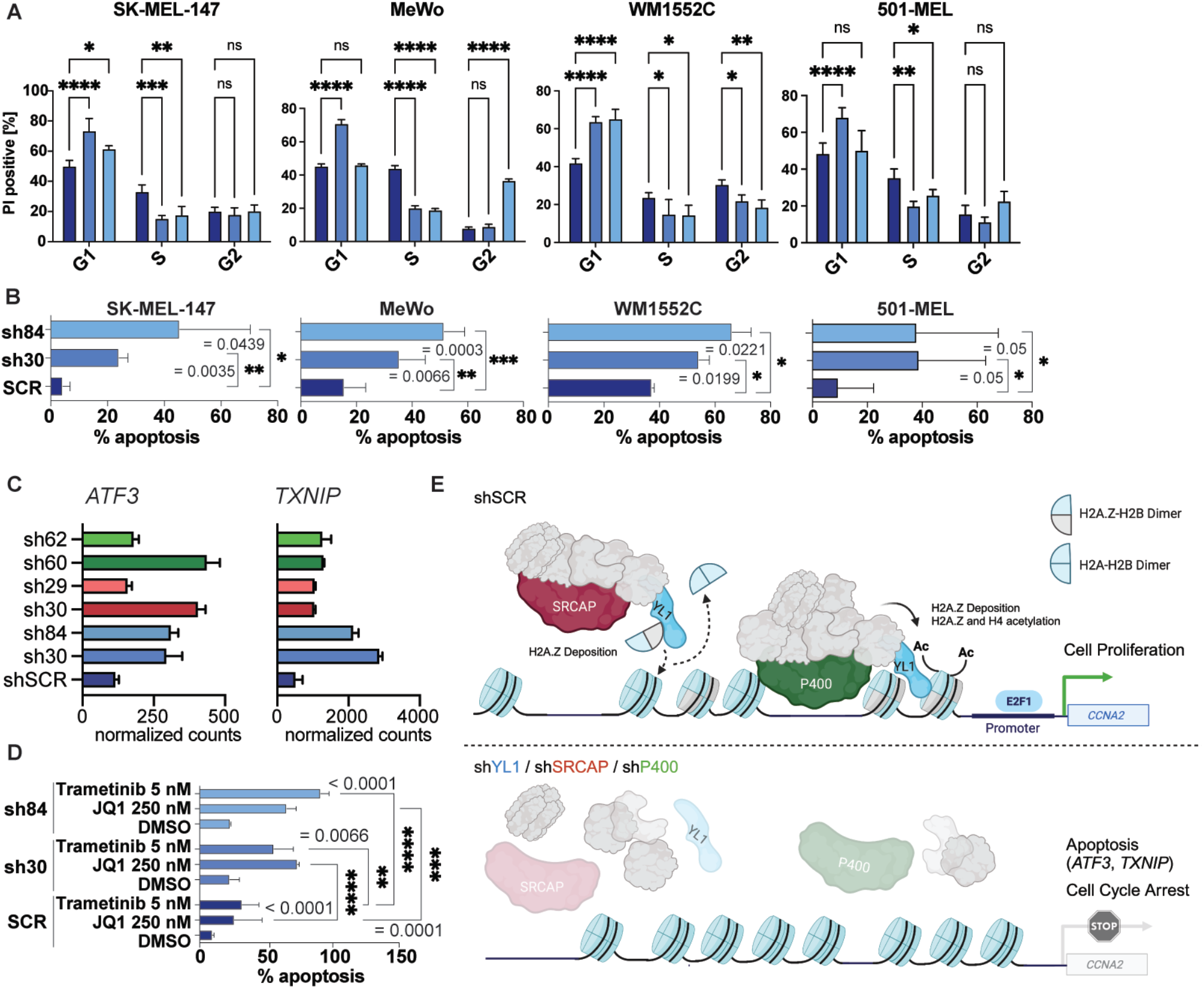
A) PI FACS analysis of YL1 knockdown cells vs. SCR controls 6 days post-infection. Significance calculated using one-way ANOVA. B) Annexin V FACS analysis of YL1 knockdown cells vs. SCR controls 6 days post-infection. Significance calculated using one-way ANOVA. C) *ATF3* and *TXNIP* mRNA expression in SCR control (dark blue), YL1 (blue), SRCAP (red) and P400 (green) knockdown cells as measured by RNA-seq. D) Annexin V FACS analysis of YL1 knockdown cells vs. SCR control when treated with 5 nM Trametinib or 250 nM JQ1 for 3 days starting 2 days post-infection with shRNAs. DMSO serves as solvent control. Statistical significance was calculated using two-way ANOVA. E) Working model of how H2A.Z chaperone subunits regulate cell cycle genes.

We therefore next aimed to identify regulators of apoptosis or cell death pathways that were deregulated in YL1 KD, but not H2A.Z.1 or H2A.Z.2 KD samples. We performed RNA-seq analysis upon H2A.Z.1 and H2A.Z.2 KD and overlapped DEGs with shYL1 DEGs (**Supp. Table S1**). Multiple inducers of a cellular stress response and apoptosis were found upregulated (*ATF3, TXNIP, SAT1, SATB1, BIK*) and one inhibitor of apoptosis (*KRT18*) was found downregulated in YL1 KD but not H2A.Z.1 or H2A.Z.2 KD samples. Of these, *ATF3* and *TXNIP* were highly expressed and showed the strongest upregulation upon YL1 KD but remained unchanged in H2A.Z.1 and H2A.Z.2 KD cells (**Fig. 7C**, **Supp.** Fig. 8A). Further, *ATF3* and *TXNIP* were also upregulated in SRCAP and P400 KD samples (**Fig. 7C**), which showed a proliferation defect comparable to the one of YL1 KD cells (**Supp.** Fig. 3A). Of note, *ATF3, TXNIP, SAT1, SATB1* and *BIK* were already upregulated 3 days post infection with at least one of two YL1 shRNAs, of which *ATF3* showed the strongest induction (**Supp.** Fig. 8B). Thus, in contrast to H2A.Z KD, YL1 loss does not only inhibit cell cycle progression but also induces apoptosis, which may be mediated by activation of the key stress response genes, such as *ATF3* and *TXNIP*.

Finally, we identified synergy between YL1 KD and treatment of melanoma cells with the BET inhibitor JQ1 or the MEK inhibitor Trametinib (**Fig. 7D**), which may be relevant for applications in a clinical setting. We therefore inquired whether melanocytes as healthy control cells would similarly be negatively affected by YL1 loss. Like melanoma cells, melanocytes showed induction of G1 arrest in one of two shRNAs (sh30), but no apoptosis was observed (**Supp.** Fig. 8C, D). Together, these findings highlight that the loss of the H2A.Z chaperone subunit YL1, but not H2A.Z itself, could be an effective approach in targeting melanoma cells.

## Discussion

Histone variants and their dedicated chaperones have emerged as key players in cancer initiation and progression. Remarkably, the H2A.Z histone chaperone complex SRCAP exhibits one of the highest mutational burdens among chromatin-modifying complexes across multiple cancers after the SWI/SNF complex (Chen et al., 2016). Interestingly, truncating mutations in SRCAP cause Floating-Harbor-Syndrome, a disease that manifests in growth deficiency, intellectual disability and craniofacial abnormalities and that arises from developmental defects in the neural crest lineage (Greenberg et al. 2019), the lineage of origin of melanoma (Goding 2000). More recently, mutations in the SRCAP members GAS41 and ZNHIT1 have shown to predispose women to uterine leiomyomas (Berta et al. 2021) and SRCAP mutations provide a selective advantage to human leukemia cells treated with chemotherapy via disruption of H2A.Z deposition and increased DNA repair (Chen et al. 2023). Moreover, different components of the SRCAP or P400-TIP60 complexes, including the SRCAP helicase, YL1, GAS41, RUVBL1 and RUVBL2 were shown to be upregulated in cancer (Ghiraldini et al. 2021). SRCAP expression is elevated in approximately 60% of colon cancers (Moon et al. 2021) and drives androgen-dependent cell growth of prostate cancer (Slupianek et al. 2010). In fact, depletion of the SRCAP and P400-TIP60 shared subunit GAS41, which contains a lysine acetyl reader (YEATS) domain, suppresses growth and survival of lung cancer cells via impaired H2A.Z deposition (Hsu et al. 2018a). Here, we focused on the role of H2A.Z chaperone complexes in melanoma via deposition of its substrate H2A.Z into chromatin. To our knowledge, the role of mutations or misexpression of SRCAP or P400-TIP60 subunits in the context of melanoma has remained elusive.

In this study, we demonstrate that the H2A.Z chaperone subunits YL1, SRCAP and P400 interact to a similar degree with both H2A.Z.1 and H2A.Z.2 variants in melanoma cells. Of note, while H2A.Z.1 and H2A.Z.2 have similar genomic localization (Vardabasso et al. 2015; Greenberg et al. 2019), they may have specific interactors, allowing them to regulate both distinct and overlapping sets of genes in a context-dependent manner (Lamaa et al. 2020). Importantly, in melanoma cells neither SRCAP nor P400 were able to compensate for the loss of the other subunit in depositing H2A.Z. Furthermore, we demonstrated that YL1 and the SRCAP-specific subunit ZNHIT1 co-localize with H2A.Z in melanoma chromatin at active promoter regions that are functionally linked to cell cycle regulation and mitosis. We and others have shown that H2A.Z isoforms interact with BRD2 and that they co-localize at active promoters (Vardabasso et al. 2015; Draker et al. 2012). We also found a large proportion of H2A.Z peaks with low signal for YL1 and ZNHIT1, which showed features of active or inactive enhancers. Studies in mouse embryonic stem cells have demonstrated that H2A.Z is incorporated into bivalent chromatin regions via Srcap and p400-Tip60, and that its monoubiquitylation antagonizes Brd2 binding (Surface et al. 2016; Hsu et al. 2018b). Thus, we expect that the inactive enhancer regions we identified in melanoma cells may contain H2A.Zub.

We found that targeting YL1, SRCAP or P400 subunits most dramatically affected the expression of genes with a strong H2A.Z peak in their promoter that were cell cycle or P53 pathway associated. While we can’t exclude the possibility that promoter-bound cell cycle or P53 genes may additionally be regulated by H2A.Z-bound enhancers (i.e., Cluster 1 regions), we focused our studies on promoter-driven effects, due to the striking co-localization of H2A.Z and both chaperone subunits YL1 and ZNHIT1 at those sites (Cluster 2). For example, a significant upregulation was observed for P53 pathway genes such as *CDKN1A*, *TXNIP* and *BAX*, whose promoters were bound by H2A.Z, YL1 and ZNHIT1 and thus identified as direct H2A.Z-YL1 targets. H2A.Z-mediated repression of stress-induced genes has been described (Lindstrom et al. 2006), specifically of the p53 downstream effector *p21* (*CDKN1A*) (Gévry et al. 2007b). Recently, Sun et al. reported that BRD8, a member of the P400-TIP60 complex, sequesters H2A.Z to p53 target loci causing a repressive chromatin state (Sun et al. 2023). How H2A.Z fosters a repressive chromatin state at these loci remains largely unexplored but is possibly linked to its PTMs and/or interactors.

Besides induction of P53 pathway genes following YL1, SRCAP and P400 KD we observed a downregulation of *E2F1* as well as other key mediators of the E2F signature such as *CDK1* and *CCNA2*. Intriguingly, we found a large proportion of these E2F1 target genes to be under control of YL1-dependent H4 acetylation at their promoter region, including *E2F1*, *CCNA2*, *BARD1* and *CDK1*. Thus, H2A.Z chaperones may support expression of these genes not only by deposition of H2A.Z, but also by acetylation of histone H4, fostering an open and active chromatin structure. H4 acetylation is likely driven by the P400-TIP60 complex that can acetylate both H2A and H4 histone tails (Altaf et al. 2010; García-González et al. 2020). The regions of increased H4ac following YL1 KD remain largely unexplored, as they were not bound by H2A.Z or its chaperones.

Together, our data emphasizes the role of H2A.Z and its chaperones in suppressing P53 pathway genes, while driving E2F1-dependent gene expression, and consequently, cell cycle regulation in melanoma. Since E2Fs play a major role in driving melanoma malignancy, especially in BRAF-resistant tumors (Liu et al. 2019), targeting H2A.Z chaperone subunits may be of therapeutic relevance in recurrent or treatment-resistant melanoma cases. Here we demonstrated that the YL1 subunit is highly expressed in melanoma cell lines and primary melanoma patient samples and speculate that its interaction with H2A.Z could be targeted by small molecules. In fact, the crystal structure of the YL1 ZID in complex with the H2A.Z/H2B dimer was resolved (Latrick et al. 2016a; Liang et al. 2016). These studies provided the molecular basis and specificity of H2A.Z/H2B recognition by YL1, and showed for that YL1 is essential for the final step of H2A.Z nucleosomal deposition (Latrick et al. 2016a; Liang et al. 2016). The implications of this specific binding and whether it is druggable remain to be explored; however, targeting the interaction with YL1 may be a viable strategy to prevent H2A.Z chromatin incorporation. Future studies will need to reveal whether there is a therapeutic window of YL1 inhibition in melanoma therapy without adversely affecting healthy cells.

## Materials and Methods

### Cell Culture

Melanoma cell lines SK-MEL-147, 501-MEL, MeWo and A375 were cultured in DMEM supplemented with 10% FBS, 100 IU of penicillin and 100 μg/mL of streptomycin. SK-MEL-239 were grown in RPMI supplemented with 10% FBS, 100 IU of penicillin and 100 μg/mL of streptomycin. Primary Melanoma cell lines WM35, WM39, WM115, WM1789, WM1552c, WM1340, WM902-B, WM793 were cultured in Tumor 2% media (80% MCDB 153 media, 20% Leibovitz’s L-15 media, 2% FBS, 5 μg/mL bovine insulin, 1.68 mM CaCl2, and 100 IU of penicillin and 100 μg/mL of streptomycin). Normal human melanocytes were grown in Melanocyte Growth Media 254 supplemented with Human Melanocyte Growth Supplement-2 (Life Technologies), calcium chloride (0.3 μM), phorbol 12-myristate 13-acetate (PMA; 10 ng/mL), and antibiotic antimycotic solution (1%). For more details on cell lines, see **Table 1**.

**Table 1.** Cell lines used in this study.

| **Cell Line** | **Melanoma Type** | **Mutations** (Data from Cellosaurus (Bairoch 2018)) |
| --- | --- | --- |
| SK-MEL-147 | Metastatic | NRAS (Gln61Arg) |
| MeWo | Metastatic | CDKN2A (Arg80Ter)<br>FGFR1 (Pro252Ser)<br>MAPK3 (Pro246Ser)<br>TP53 (Gln317Ter) |
| 501-MEL | Metastatic | CDKN2A (homozygous deletion)<br>PTEN (homozygous deletion)<br>BRAF (Gly469Arg)<br>BRAF (Val600Glu) |
| SK-MEL-28 | Derived from Skin | BRAF (Val600Glu)<br>CDK4 (Arg24Cys)<br>EGFR (Pro753Ser)<br>PTEN (Thr167Ala)<br>TERT (57A>C)<br>TP53 (Leu145Arg) |
| SK-MEL-239 | Metastatic | BRAF (Val600Glu) |
| WM1552c | Primary | CDKN2A (homozygous deletion)<br>CDKN2B (homozygous deletion)<br>BRAF (Val600Glu)<br>PTEN (634+5G>T)<br>TP53 (Arg248Gln) |

### Plasmids and Infections

Lentiviral plasmids encoding shRNAs against *VPS72* (YL1), *SRCAP*, *P400, H2AFV (*H2A.Z.2*), H2AFZ* (H2A.Z.1), and *TIP60* (*KAT5*) were obtained from the TRC shRNA library and sequences are listed below (see **Table 2**). shSCR (sh_scrambled) served as control. For CRISPR-mediated knockout, gRNAs targeting *VPS72* or *SRCAP* were cloned into the lentiCRISPRv2 (addgene: #52961). For gRNA sequences, see **Table 3**. eGFP-fusion constructs of H2A, H2A.Z.1 and H2A.Z.2 were generated previously (Vardabasso et al. 2015). Virus production and infections were performed using standard procedures (Kapoor et al., 2010). In brief, 5x10^5 cells were seeded into 10cm plates and infected with shRNA virus the following day. Subsequently, cells were washed twice with PBS and selected in DMEM medium containing puromycin (2 μg/mL) for 24 hours.

**Table 2.**
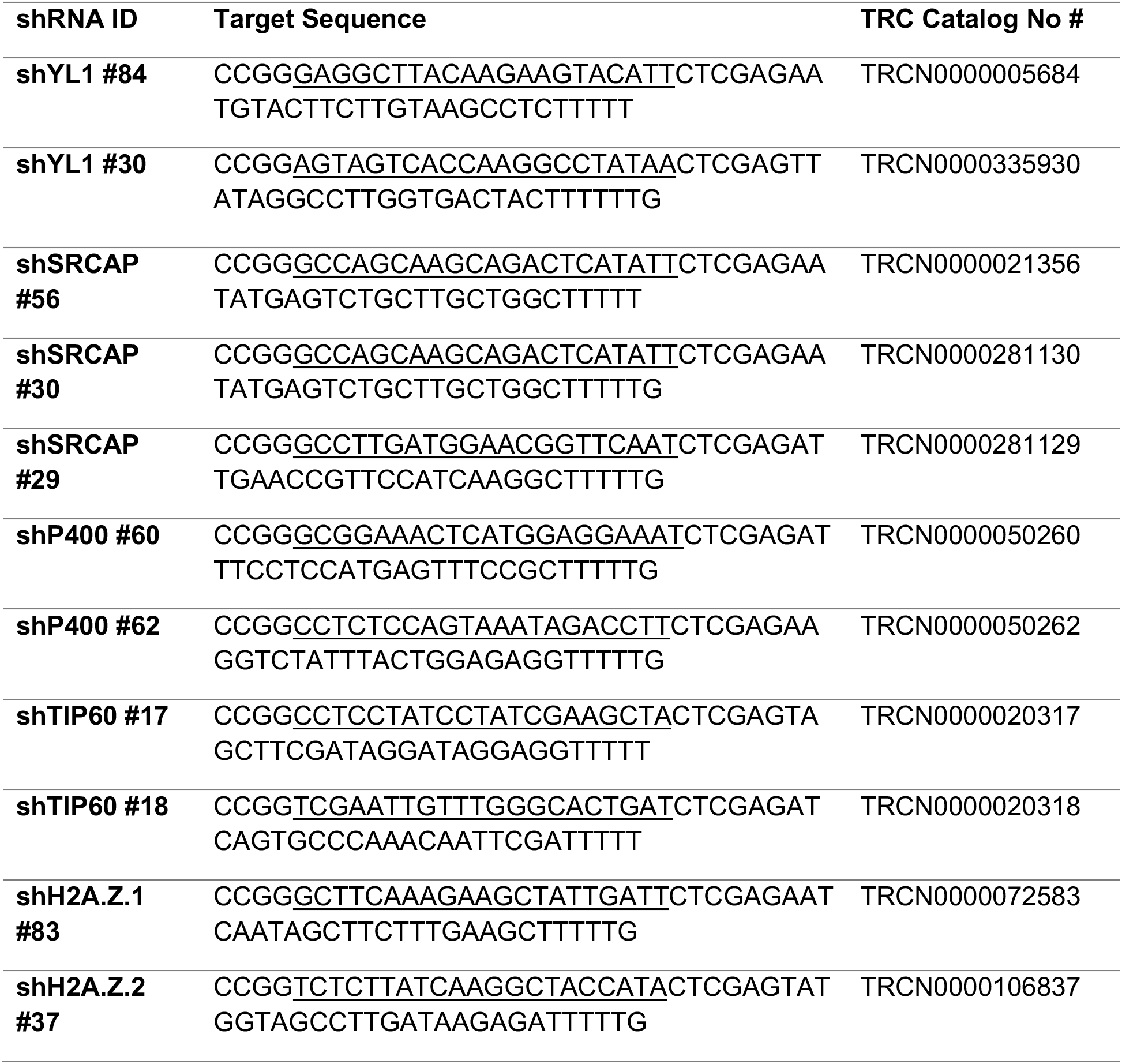
shRNAs used in this study.

**Table 3.**
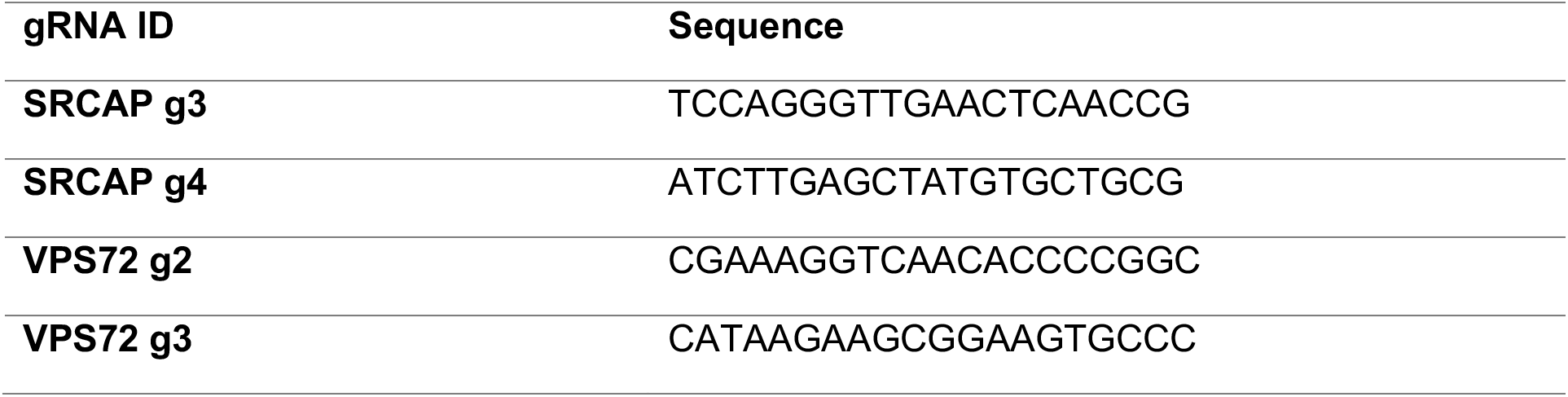
lentiCRISPRv2 guide RNAs used in this study.

### Chromatin Fractionation, Whole Cell Protein Extraction, and Immunoblotting

For chromatin extraction, cell pellets were lysed on ice for 8 min in buffer A (10 mM HEPES pH 7.9, 10 mM KCl, 1.5 mM MgCl2, 0.34 M Sucrose, 10% glycerol supplemented with protease inhibitors and 1 mM DTT) + 0.1% triton x-100. Samples were centrifuged for 5 min at 1850 g (supernatant contains cytoplasmic fraction) and pellets washed with 1 mL of buffer A (supplemented with protease inhibitors and 1 mM DTT). Samples were centrifuged for 5 min at 1850 g, pellets resuspended in No Salt Buffer (3 mM EDTA, 0.2 mM EGTA supplemented with protease inhibitors and 1 mM DTT) and kept on ice for 30 min with occasional vortexing. Samples were centrifuged for 5 min at 1850 g (supernatant contains soluble nuclear fraction) and chromatin pellets were resuspended in 200 μL buffer A (supplemented with protease inhibitors and 1:200 benzonase). Pellets were solubilized for 15 min at 37 degrees, shaking and subsequently used for Western blot analysis. For whole-cell extraction, cells were lysed on ice for 30 minutes in RIPA lysis buffer + benzonase (Millipore Sigma) (supplemented with protease inhibitors). Lysates were sonicated on high level, 5 cycles 30s ON, 30s OFF and centrifuged at 10,000 g for 10 minutes. Protein concentrations were quantified using BCA (Pierce). Lysates were mixed with 4× Laemmli loading buffer with subsequent boiling prior to immunoblotting.

### Cell Proliferation and Crystal Violet Staining

For proliferation curves, cell counts were tracked and quantified over time in the Incucyte Live-Cell Imaging System (Essen Bioscience). Following infection, cells were selected in puromycin (2 μg/mL) for 24 hours and then continuously measured for confluence in 24-hour time intervals. Non-selected cells were included as reference to determine transduction efficiency (data not shown). Cell numbers were normalized to cell counts on day 1. Crystal violet staining was performed on the last day of cell counting as follows: Cells were fixed in 100% ice-cold methanol for 10 minutes and then stained in 0.5% crystal violet in 25% methanol.

### Apoptosis and Cell Cycle Flow Cytometry

PI and Annexin-V FACS analysis were performed on day 6 post infection (melanoma cells) and day 7 post infection (melanocytes). For single-parameter apoptosis analysis, floating cells were harvested and combined with trypsinized seeded cells, washed with phosphate-buffered saline, labeled with AnnexinV-FITC in binding buffer (10 mM HEPES pH 7.4, 150 mM NaCl, 5 mM KCl, 1 mM MgCL_2_, 1.8 mM CaCl_2_), and analyzed on flow cytometry. For multi-parameter apoptosis assay, cells were collected as above and stained using propidium iodide (FITC Annexin V Apoptosis Detection Kit; BD) and APC Annexin V (BD), per the manufacturer’s protocol. For cell cycle analysis, trypsinized cells were washed and resuspended in phosphate-buffered saline, stained with propidium iodide (20 μg/mL), and analyzed on flow cytometry. FACS analyses were performed on FlowJo 6.7 software and FCS Express 7 Research software.

### RNA Extraction and RNA-seq

Total RNA was extracted using RNeasy Mini Kit (Qiagen). For qRT-PCR, reverse transcription was performed with First-strand cDNA Synthesis kit (OriGene). For RNA-seq, the quality of RNA samples was assessed on a 2100 Agilent Bioanalyzer. mRNA was then extracted from 2 ug of total RNA per sample using NEXTFLEX® Poly(A) Beads 2.0 (Perkin Elmer, Austin, Texas, USA). Libraries were prepared from mRNA samples using NEXTFLEX® Rapid Directional RNA-seq Kit 2.0 (Perkin Elmer, Austin, Texas, USA). Quality of library preparation was assessed on a 2100 Agilent Bioanalyzer. Single-end 75-bp reads were sequenced on the HiSeq2500 according to the manufacturer’s guidelines (Illumina). Reads were aligned to the human reference genome (hg19/GRCh37.p13) with STAR (Dobin et al. 2013) (version 2.6.0.c) using the parameters -- runMode alignReads --sjdbOverhang 100 --outFilterMultimapNmax 10 --outFilterMismatchNmax 10 --outFilterType BySJout --outFilterIntronMotifs RemoveNoncanonicalUnannotated. Following, featureCounts from the Rsubread (Liao et al. 2019) (version 2.4.3) R package was used to assign reads to coding genes. Assigned reads were then normalized and differentially expressed genes were identified using the R package DEseq2 (version 1.30.1) (Love et al. 2014). Genes were considered expressed if the sum of raw counts was >10 for any given gene. Differentially expressed genes were called using an adjusted p value ≤ 0.05 and log2FC ≥ 0.75 or ≤ -0.75. Principal component analysis (PCA) was generated using regularized log-transformed reads with the DEseq2 package. Heatmaps were generated with the pheatmap (version 1.0.12) package, using DEseq2 normalized counts.

### Mononucleosome Immunoprecipitation (IP)

Cells were lysed, isolated for nuclear material, and digested with MNase as described (Vardabasso et al. 2015). In brief, for each IP 8x10^7 cells were lysed in 1 ml ice-cold PBS/0.3% triton x-100 (with protease inhibitors) and incubated for 10 min on ice with occasional vortexing. Cells were then pelleted for 10 min at 1000 g, 4 degrees. Pellet was washed with PBS and resuspended in 500 μl EX-100 buffer (10 mM HEPES pH 7.6, 100 mM NaCl, MgCl_2_, 0.5 mM EGTA, 10% v/v glycerol, with protease inhibitors). Chromatin was solubilized for 20 min with MNase at 37 degrees. Reaction was stopped by adding 1/50th of 0.5M EGTA. Samples were centrifuged for 5 min at 1000 g, 4 degrees and supernatant (S1) was used for IP. For S2, pellets were resuspended in RES Buffer (PBS, 150 mM NaCl, 2 mM EDTA, 0.1% tritron x-100) and rotated at 4 degrees O/N. Samples were centrifuged for 30 min at 1000 g, 4 degrees C. Supernatant is S2. For IP, 25 μl slurry beads were equilibrated in EX100 buffer and then incubated with S1 mononucleosomes of 8x10^7^ cells for 2.5 h at 4°C (rotating). Beads were washed twice in wash-buffer 1 (10mM Tris-HCl, pH 7.5, 150mM NaCl, 1mM DTT, 1xCPI), followed by 2 washes in wash-buffer 2 (10mM Tris-HCl, pH 7.5, 150mM NaCl, 0.1% NP-40). Samples were then boiled with Laemmli buffer for immunoblot analysis.

### Clinical Specimens

Formalin-fixed paraffin-embedded human nevi and melanoma tumor resections and clinical outcomes were obtained from the Icahn School of Medicine at Mount Sinai Department of Dermatology and Pathology and the Mount Sinai Biorepository with approval from the Institutional Review Board at Mount Sinai (IRB project number 16-00325).

### Immunohistochemistry

Formalin-fixed and paraffin-embedded clinical specimens sectioned at 3 or 5-μm were baked at 60°C for 1 hour and deparaffinized in graded xylene and ethanol washes. Antigen retrieval was performed in citrate-based buffer (10mM sodium citrate, 0.05% Tween 20, pH 6.0) in heated water for 10 minutes. Samples were soaked in 3% hydrogen peroxide, blocked with 2% horse serum (in 1% BSA, 0.1% Triton X-100, 0.05% Tween-20, and 0.05%) for 30 minutes, and incubated overnight with anti-YL1 (1:400; Abcam ab72506) prepared in blocking buffer. Slides were developed in ImPRESS HRP anti-mouse/rabbit IgG (Vector) as the secondary, ImmPACT NovaRed as the chromogen, and Mayer’s hematoxylin (Volu Sol) for counterstaining. Slides were washed in 1% acetic acid and 0.1% sodium bicarbonate prior to dehydration in graded ethanol and xylene, prior to mounting with Permount (Sigma SP15-100). Slides were stained with H3 (1:300, Abcam ab1791) positive control for assessment of tissue quality. Slides were scored by 2 independent dermatopathologists in a blinded fashion using a 4-point scale in terms of number of cells stained (1=0-25% positive cells; 2=25-50% positive cells; 3=50-75% positive cells; 4=75-100% positive cells) and staining intensity (1 = absent, 2 = weak, 3 = moderate, 4 = strong) (**Supp. Table S4**). The 2 scores are multiplied to yield a single score per pathologist, and subsequently averaged together to yield 1 score per slide.

### ChIP-sequencing

For YL1 and H4ac ChIP, SK-MEL-147 cells were (1x10-cm plate per sample) cross-linked with 1% Formaldehyde for 10 min at room temperature. For ZNHIT1 ChIP, SK-MEL-147 cells were (1x10-cm plate per sample) were double cross-linked with 0.25 M disuccinimidyl glutarate (DSG) for 45 min, followed with 1% formaldehyde for 10 min. Single and double cross-linked cells were quenched with 0.125 M glycine for 5 min at room temperature, washed 3 times in PBS and then collected in 1 mL ice-cold PBS. Chromatin was then pelleted at 1100RPM @4C for 3min and stored at -80 degrees celsius until ready for ChIP. ChIP and library preparation were performed as described (Carcamo et al. 2022). For antibody details, see **Table 4**. Libraries were sequenced on Illumina Hi-Seq2500 (75bp single-end reads).

**Table 4.** Antibodies used in this study.

| **Antibody** | **Catalog #** | **Dilution** |
| --- | --- | --- |
| **H4K5ac** | ab51997 | 1 : 1000 (WB) |
| **H4K12ac** | ab46983 | 1 : 1000 (WB) |
| **H4K16ac** | ab109463 | 1 : 1000 (WB) |
| **H4ac** | 06-866 | 1 : 2000 (WB), 5 µg (ChIP) |
| **H2A.Z** | ab4174 | 1 : 1000 (WB) |
| **H2A.Z** | PA5-21923 | 1 : 1000 (WB) |
| **H2A.Z K4ac** | ab214725 | 1 : 1000 (WB) |
| **H2A.Z K7ac** | H2A.Z K7ac | 1 : 1000 (WB) |
| **H3** | ab1791 | 1 : 2000 (WB) |
| **ZNHIT1** | ab238125 | 1 : 2000 (WB), 5 µg (ChIP) |
| **GAS41** | sc-393708 | 1 : 1000 (WB) |
| **GFP** | 1181460001 | 1 : 1000 (WB) |
| **GAPDH** | sc-32233 | 1 : 10,000 (WB) |
| **YL1** | ab112055 | 1 : 2000 (WB) |
| **P53** | sc-126 | 1 : 1000 (WB) |
| **E2F1** | 32-1400 | 1 : 100 (WB) |
| **LAMIN** | SAB4200236 | 1 : 5000 (WB) |
| **RNA Pol II (phospho S2)** | ab5095 | 1 : 2000 |
| **RNA Pol II (phospho S5)** | A304-408A | 1 : 4000 |
| **P400** | A300-541A | 1 : 1000 (WB) |
| **SRCAP** | PA5-56012 | 1 : 1000 (WB) |
| **TRRAP** | Tora Lab (IGBMC) | 1 : 500 (WB) |

### ChIP Alignment and Peak Calling

ChIP reads were aligned to the human reference genome GRCh37/hg19 using Bowtie (version 1.1.2) (Langmead et al. 2009) with parameters –l 50 –n 2 –S --best –k 1 –m 1 for ZNHIT1 and YL1 or –l 65 –n 2 –best –k 1 –m 1 for H4ac. Read quality was assessed using fastQC (Andrews 2010) (version 0.11.7). Duplicate reads were removed with PICARD (version 2.2.4) (Broad Institute). Binary alignment map (BAM) files were generated with samtools v1.9 (Li et al., 2009), and were used in downstream analysis. Significant peaks were identified using MACS2 (version 2.1.0) (Zhang et al. 2008) where q-value cut-offs were determined post-hoc, testing several q-values based on signal to background ratio. YL1 and ZNHIT1 peaks were called against matching input control with parameters --nomodel –s 75 --keep-dup 2 -q 0.005 or -q 0.05 (for ZHNIT1 ChIP). For the H4ac ChIP-seq the bam files of 2 control and 2 KD samples (shSCR 2x, and YL1 sh30 and YL1 sh84) were concatenated using samtools merge to generate ‘master’ bam files. Significant peaks were called on ‘master’ bam files and matching input controls using MACS2 for narrow peaks with -q 1e-10. Peaks in ENCODE blacklisted regions were removed. Coverage tracks were generated from BAM files for master bam files and individual replicates and conditions using deepTools (version 3.2.1) bamCoverage (Ramírez et al. 2014) with parameters --normalizeUsingRPKM --binsize 10. H2A.Z ChIP-seq in SK-MEL-147 was downloaded from previously published dataset (GSM1665991) (Vardabasso et al. 2015) and bed files were further filtered to retain peaks with better enrichment (peaks with 50 read counts or fewer were excluded, as quantified by the subread featureCounts function). ChIP-seq enrichment plots were visualized on the IGV genome browser (Robinson et al. 2011). Enhancers and super-enhancers in SK-MEL-147 cells were identified by ROSE (Whyte et al. 2013; Lovén et al. 2013) using previously published H3K27ac ChIP-seq data (Carcamo et al. 2022).

### Cluster definitions

Clusters were defined based on the differential and shared occupancy of H2AZ, YL1 and ZNHIT1. Venn diagrams and bed files of the different genomic regions were generated using the Intervene (v0.6.4) package. Cluster 1 regions (n = 24427) correspond to significant regions exclusive to H2AZ, Cluster 2 regions (n = 6324) correspond to significant regions shared between H2AZ, YL1 and ZNHIT1, Cluster 3 regions (n = 5146) correspond to significant regions exclusive to YL1, Cluster 4 regions (n = 1765) correspond to significant regions exclusive to ZNHIT1, and Cluster 5 regions (n = 1873) correspond to significant regions exclusive to YL1 and ZNHIT1.

### Metagenes and heatmaps

Metagene and heatmaps of genomic regions were generated with deepTools (version 3.2.1) (Ramírez et al. 2014). The command computeMatrix was used to calculate scores at genomic regions and generate a matrix file to use with plotHeamap or plotProfile, to generate heatmaps or metagene profile plots, respectively.

### Differential H4ac analysis

The H4ac ChIPseq BAM files of all the conditions (shSCR 2x and shYL1 (sh30 and sh84)) were combined into a single BAM file and significant peaks were called using MACS2 as described above to generate a universe of regions present in all conditions. Regions within 500 bases were merged with bedtools merge to better capture the ChIP-seq enrichment signal. Following, Diffbind (version 3.4.11) (Stark and Brown; Ross-Innes et al. 2012) was used to generate PCA plots and to quantify the reads in the universe of regions, normalize counts and estimate significantly differential enriched peaks with default parameters (normalize=DBA_NORM_LIB, library=DBA_LIBSIZE_FULL, method=DBA_DESEQ2). Significant differentially enriched regions were called using an adjusted p-value < 0.05 (using the Benjamini and Hochberg procedure).

### Genomic annotation analysis

Promoters (-1 kb to +1 kb) relative to the TSS were defined according to the human GRCh37/hg19 Gencode v19 genome annotation. Promoters of expressed genes were classified as active promoters whereas all other promoters were defined as weak/inactive promoters. The ChIPSeeker (version 1.26.2) (Yu et al. 2015) package was modified and used to determine feature distribution for peak sets. Enhancers identified by ROSE were defined as “active enhancers”, whereas all other distal regions were defined as weak/poised enhancers.

## Data Availability

RNA-seq data is published in GSE242227.

ChIP-seq data is published in GSE246121.

## Acknowledgements

We thank the Bernstein lab, especially Elena Grossi for her valuable feedback and contributions; Theodora Smith for her technical assistance; Kunal Kumar, Roberto Sanchez, Robert Devita (ISMMS) and Michael Keogh (Epicypher) for their expertise and advice, Didier Devys and Laszlo Tora (IGBMC), Xiaobing Shi (VAI) and David Dominguez-Sola (ISMMS) for sharing reagents. The authors acknowledge the Center for Advanced Genomics Technology, the Dean’s Flow Cytometry Core and the Biorepository and Pathology Core at ISMMS. This work was also supported by the Bioinformatics for Next Generation Sequencing (BiNGS) shared resource facility within the Tisch Cancer Institute at ISMMS, which is partially supported by NIH grant P30CA196521. This work was also partially supported by Scientific Computing at ISMMS and supported by the Clinical and Translational Science Awards (CTSA) grant UL1TR004419 from the National Center for Advancing Translational Sciences as well the Office of Research Infrastructure of the NIH under award number S10OD026880. This work was supported by DFG [429315233] to S.J., American Skin Association Research Grant for Skin Cancer and Melanoma to C.V., HHMI Medical Research Fellows Award to J.D. and NCI/NIH [R01CA154683] to E.B.

## Author Contributions

S.J., C.V. and E.B. conceived of this study. S.J., C.V., and J.D. designed and performed experiments and interpreted data. SC analysed data, R.S. and R.P. scored IHC samples, A.M. performed experiments. D.H. guided experiments and data analysis. E.B. supervised the project and interpreted data. S.J. and E.B. wrote the manuscript with input from other authors

