## Supplementary Figures for "H2A.Z chaperones converge on histone H4 acetylation for melanoma cell proliferation"

**Jostes et al., Supp. Fig. 1**

A

| Welch Difference | Z.1 vs. H2A | Z.2 vs. H2A |
|---|---|---|
| YL1 | 5.40 | 5.25 |
| SRCAP | 3.10 | 3.38 |
| P400 | 1.21 | 1.33 |

B

| | | H2A.Z.1 vs H2A | H2A.Z.2 vs H2A |
|---|---|---|---|
| P400-TIP60 | TRRAP | + | + |
| | EP400 | + | + |
| | EPC1 | - | - |
| | EPC2 | - | - |
| | BRD8 | | |
| | KAT5 / TIP60 | | |
| | ING3 | | |
| | MBDT1 | | |
| | MEAF6 | - | - |
| | MRGP1 | | |
| Common | DMAP1 | ++ | ++ |
| | GAS41 | +++ | +++ |
| | ACTL6A | - | - |
| | ACTIN | | |
| | YL1 | +++ | +++ |
| | RUVBL1 | + | + |
| | RUVBL2 | ++ | ++ |
| SRCAP | SRCAP | +++ | +++ |
| | ACTR6 | +++ | +++ |
| | ZNHIT1 | ++ | ++ |

| | |
|---|---|
| + | 1-2 |
| ++ | 2-3 |
| +++ | >3 |
| - | not enriched |
| (grey) | not detected |

C

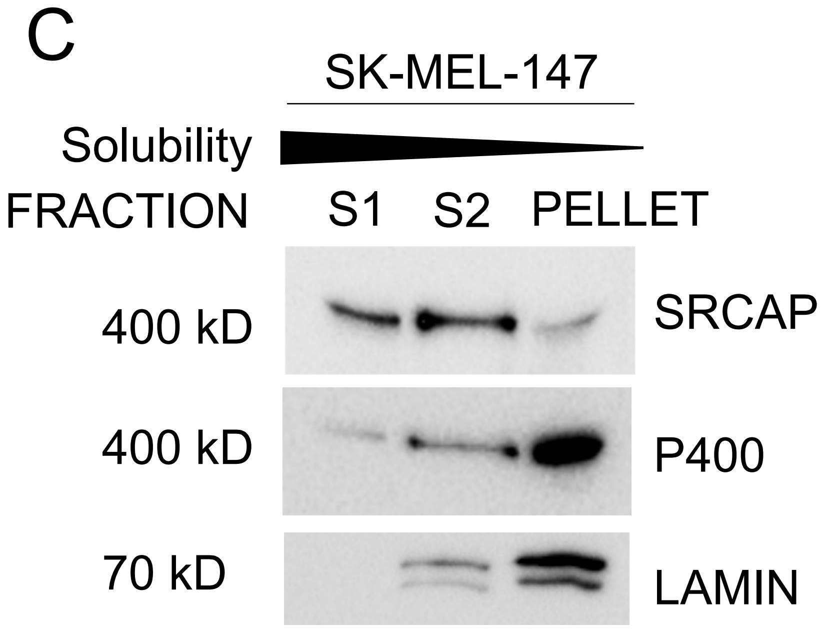


D

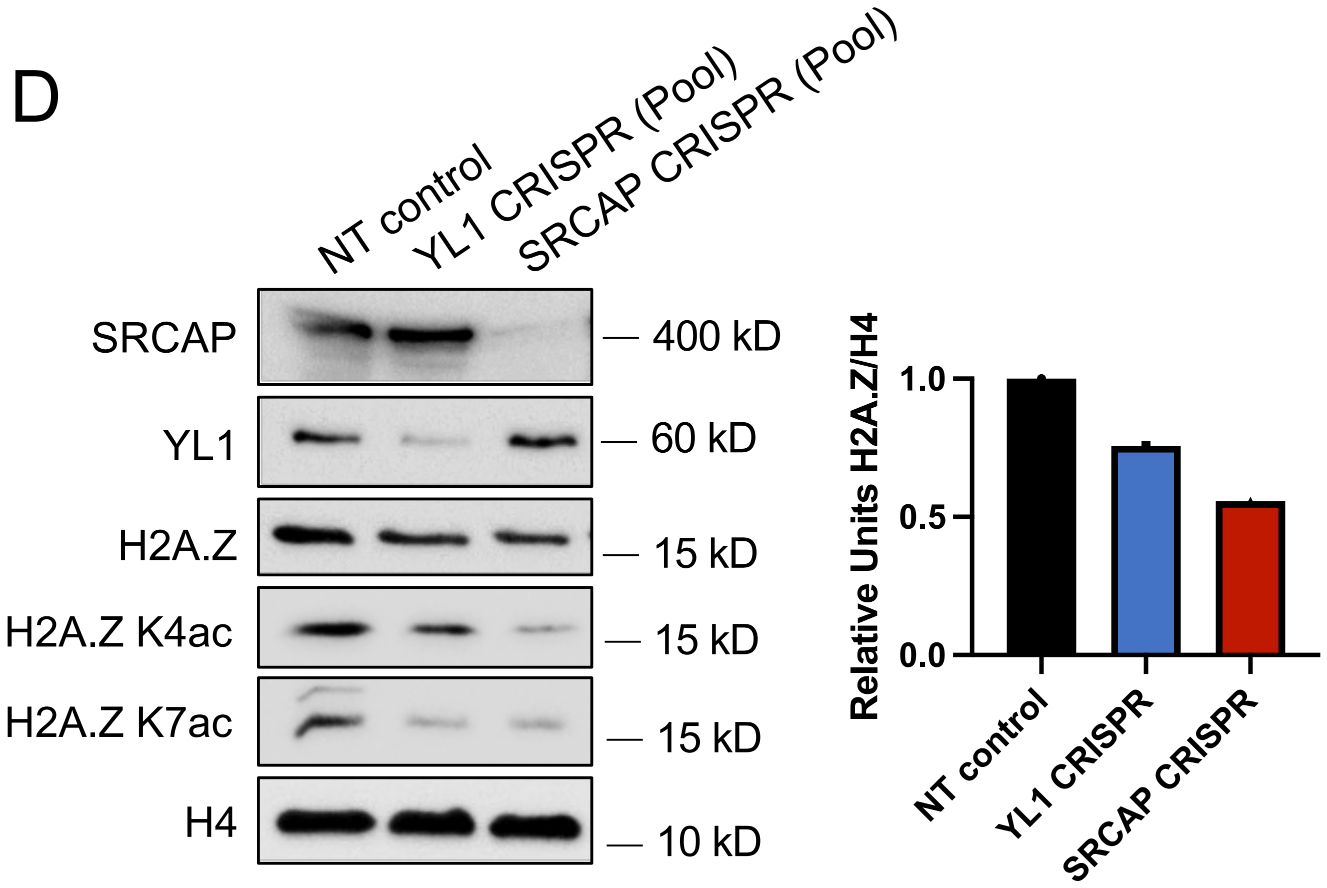


**Supp. Figure 1:** A, B) Mass Spectrometry results (published in Vardabasso et al. (2015)) of anti-GFP co-IP in SK-MEL-147 ($NRAS^{Q61R}$) cells expressing GFP-H2A, GFP-H2A.Z.1 or GFP-H2A.Z.2. Welch t-test differences shown for H2A.Z.1 or H2A.Z.2 over H2A. C) Western blots of MNase-digested chromatin extracts of SK-MEL-147 cells (S1 = readily soluble fraction, S2 = solubilized after O/N rotation in RES buffer, Pellet = insoluble fraction; see materials and methods for details). D) Western blots of H2A.Z and H2A.Z chaperone subunits in SK-MEL-147 chromatin lysates after CRISPR-mediated knockout of *VPS72* (YL1) and *SRCAP*. Cells were analyzed as a pool (as a mixed population of YL1 / SRCAP -/-, -/+, +/+). H4 serves as loading control. Bar graph (right) shows quantification of H2A.Z levels relative to H4 loading control.

**Jostes et al., Supp. Fig. 2**

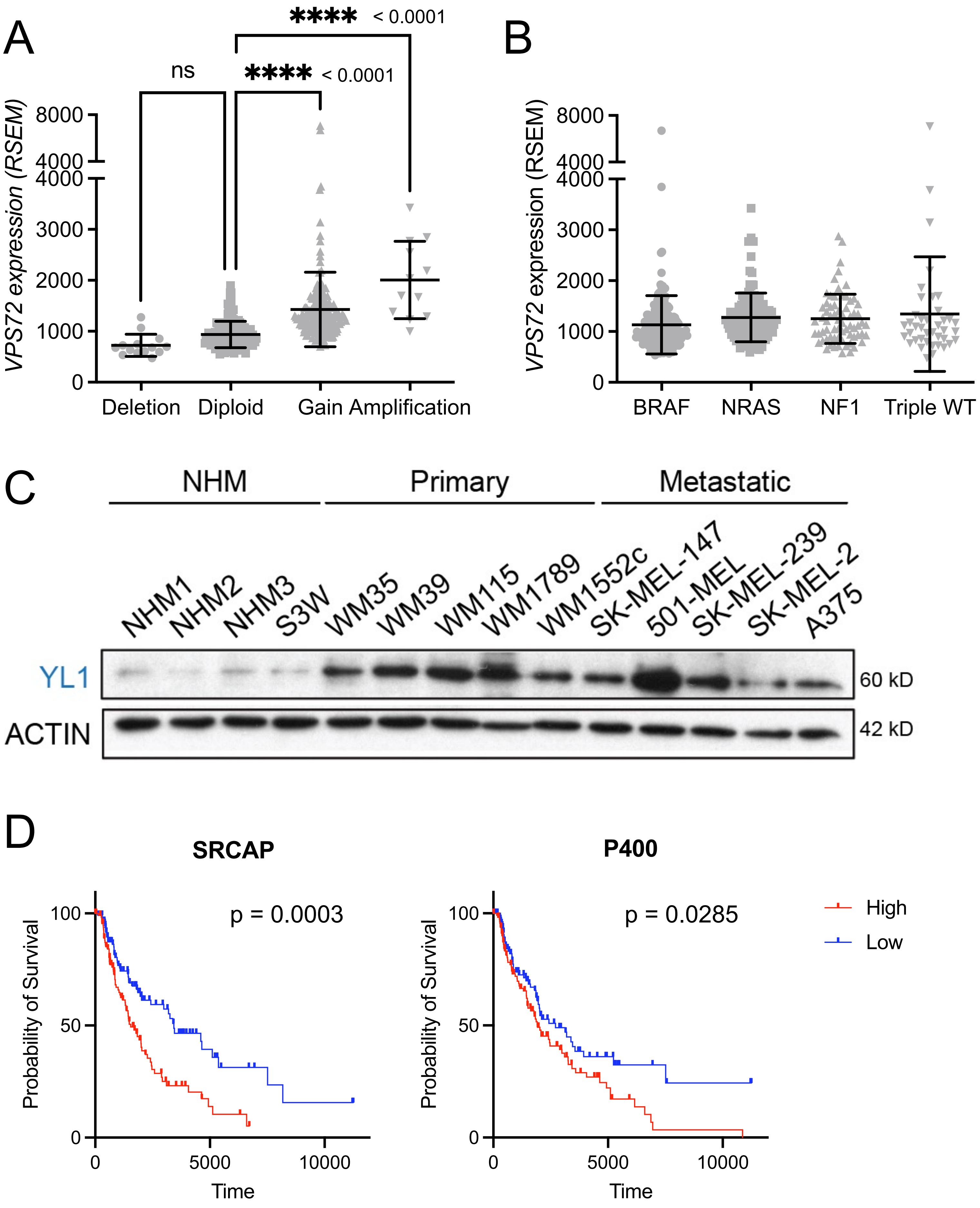


**Supp. Figure 2:** A) *VPS72* expression in Skin Cutaneous Melanoma (TCGA, PanCancer Atlas) stratified by copy number data (Deletion n=14, Diploid n=153, Gain n=188, Amplification n=12). Error bars indicate mean and SD. B) *VPS72* expression in Skin Cutaneous Melanoma (TCGA, PanCancer Atlas) stratified by melanoma subtype (BRAF n=193, NRAS n=115, NF1 n=68, Triple WT n=40). Error bars indicate mean and SD. C) YL1 protein levels in whole cell lysate of normal human melanocytes (NHM), primary melanoma, and metastatic melanoma cell lines. ACTIN served as loading control. D) Survival of patients with high vs. low *SRCAP* and *EP400* expression in TCGA melanoma cohort of primary and metastatic melanoma (n=228). Significance calculated with logrank Mantel-Cox test.

Jostes et al., Supp. Fig. 3

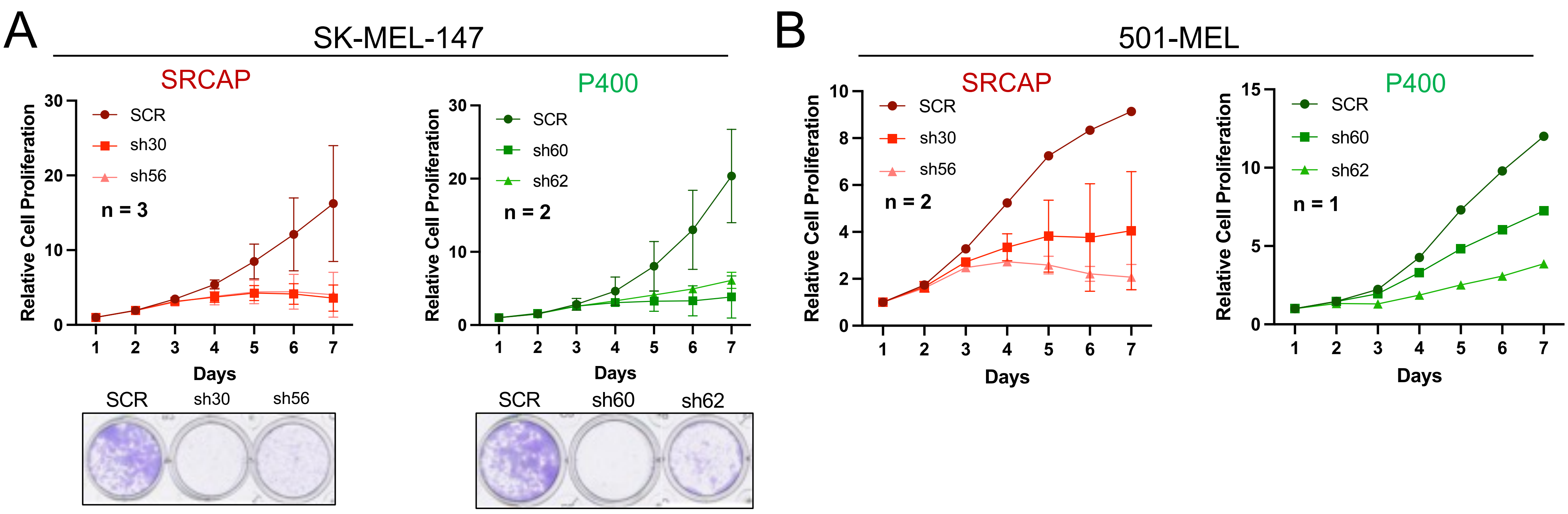


**Supp. Figure 3:** A, B) Proliferation of SK-MEL-147 and 501-MEL melanoma cells after SRCAP (red) or P400 (green) knockdown over the course of 7 days. Error bars indicate mean and SD. Number of biological replicates shown for each knockdown experiment. In (A) crystal violet staining indicates cell density of SK-MEL-147 cells at 7 days post-knockdown.

Jostes et al., Supp. Fig. 4

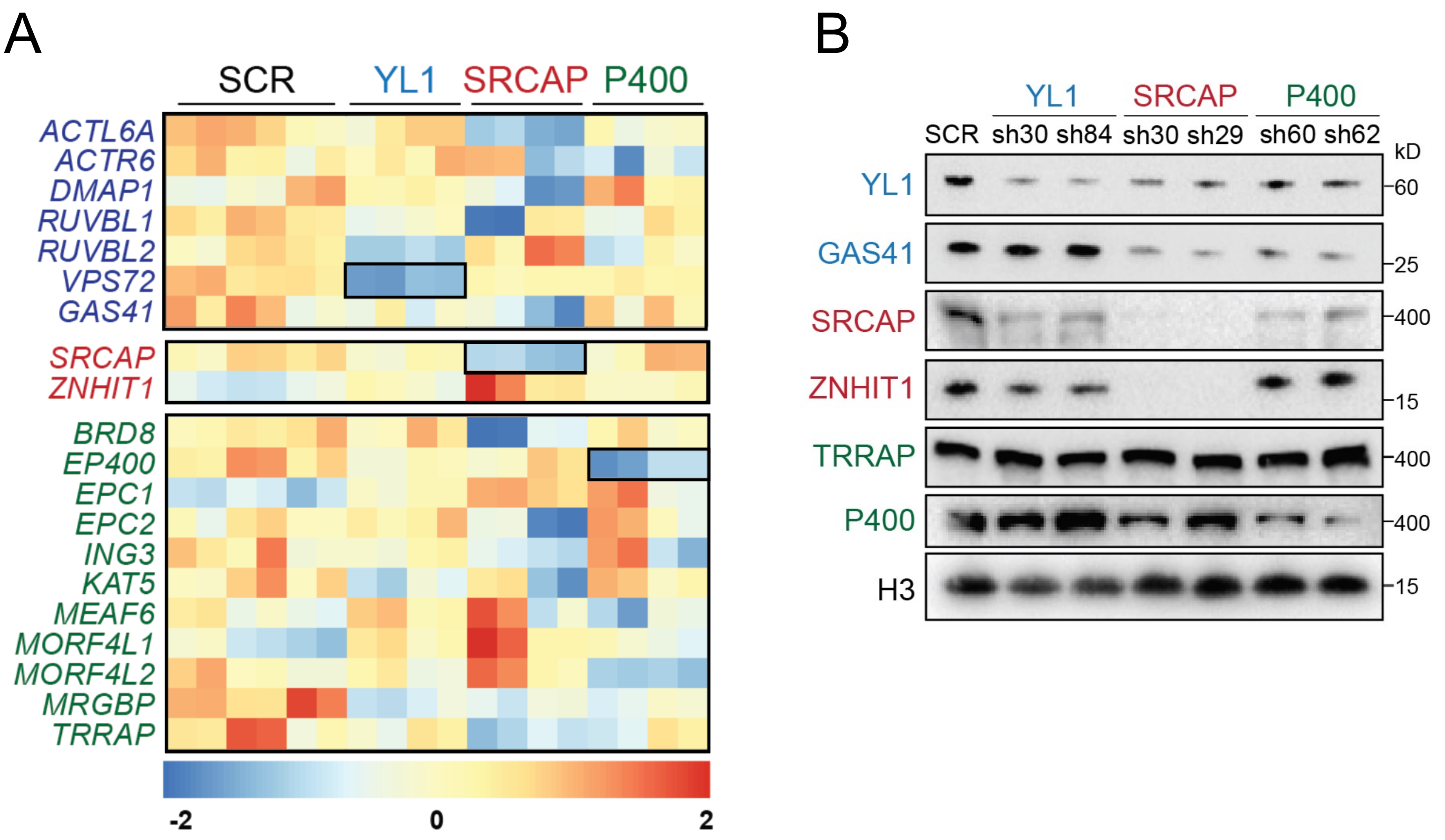


**Supp. Figure 4:** A) Heatmap showing mRNA expression levels of SRCAP (red), P400 (green) and shared (blue) chaperone subunits in SK-MEL-147 cells 6 days post infection with YL1, SRCAP or P400 shRNAs or SCR controls. B) Western blots of SRCAP (red), P400 (green) and shared (blue) chaperone subunits in chromatin lysates of YL1, SRCAP and P400 knockdown samples compared to SCR control. H3 serves as loading control.

**Jostes et al., Supp. Fig. 5**

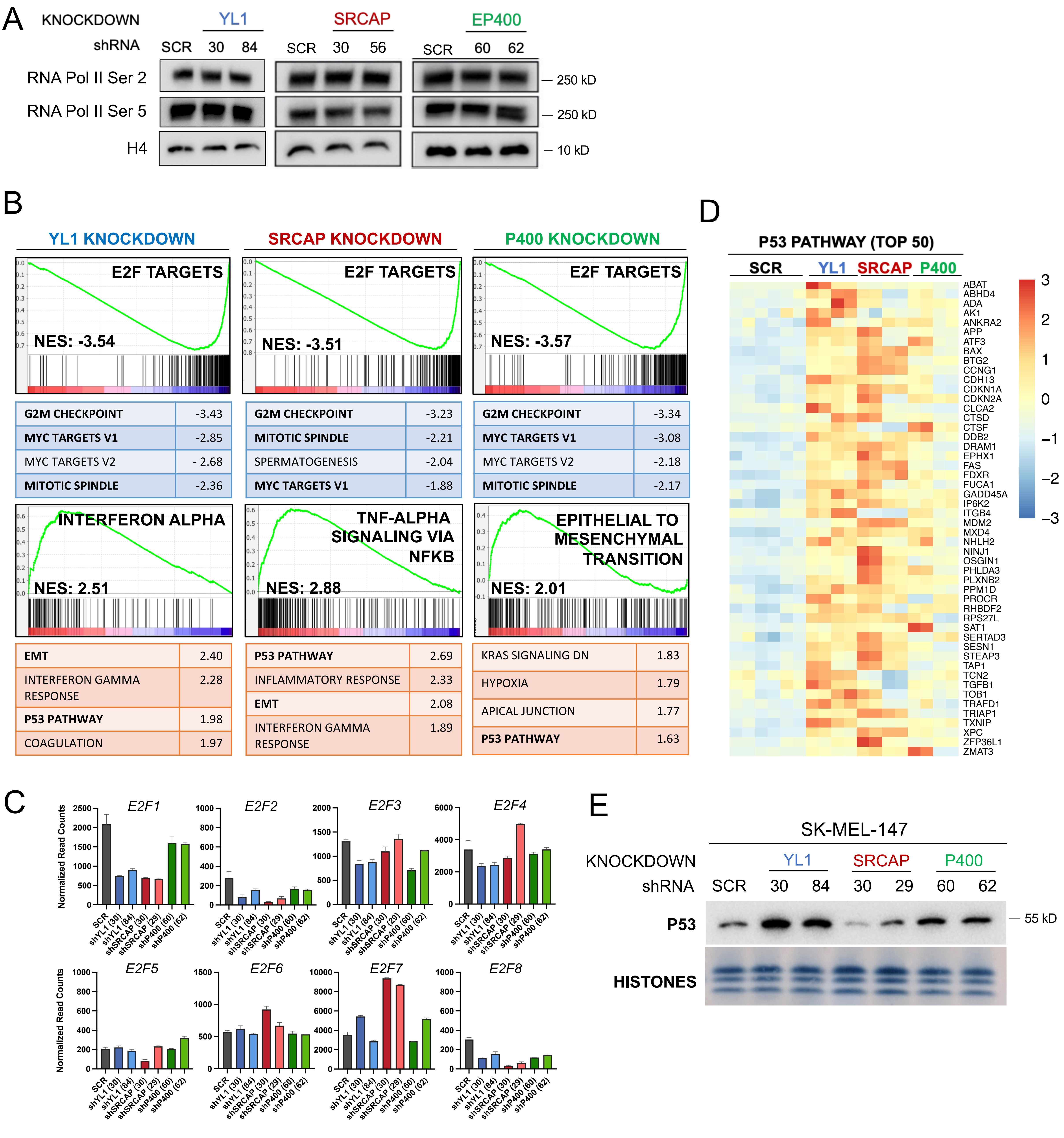


**Supp. Figure 5:** A) RNA Pol II Ser2 and Ser5 phosphorylation levels in chromatin lysates of SK-MEL-147 cells following YL1, SRCAP and P400 knockdown. H4 serves as loading control. B) Gene Set Enrichment Analysis (GSEA) of genes down- (blue shading) and up-regulated (red shading) after YL1, SRCAP and P400 knockdown. Values indicate normalized enrichment score (NES). C) Expression levels of E2F family members in YL1, SRCAP and P400 knockdown cells and SCR controls as measured by RNA-seq. Error bars indicate mean with SD. D) Heatmap showing normalized counts of top 50 upregulated P53 pathway genes (as identified by GSEA) in YL1, SRCAP and P400 knockdown samples compared to SCR controls. E) Western blot of P53 levels in YL1 and P400 knockdown cells. Amido black staining of histones serves as loading control.

**Jostes et al., Supp. Fig. 6**

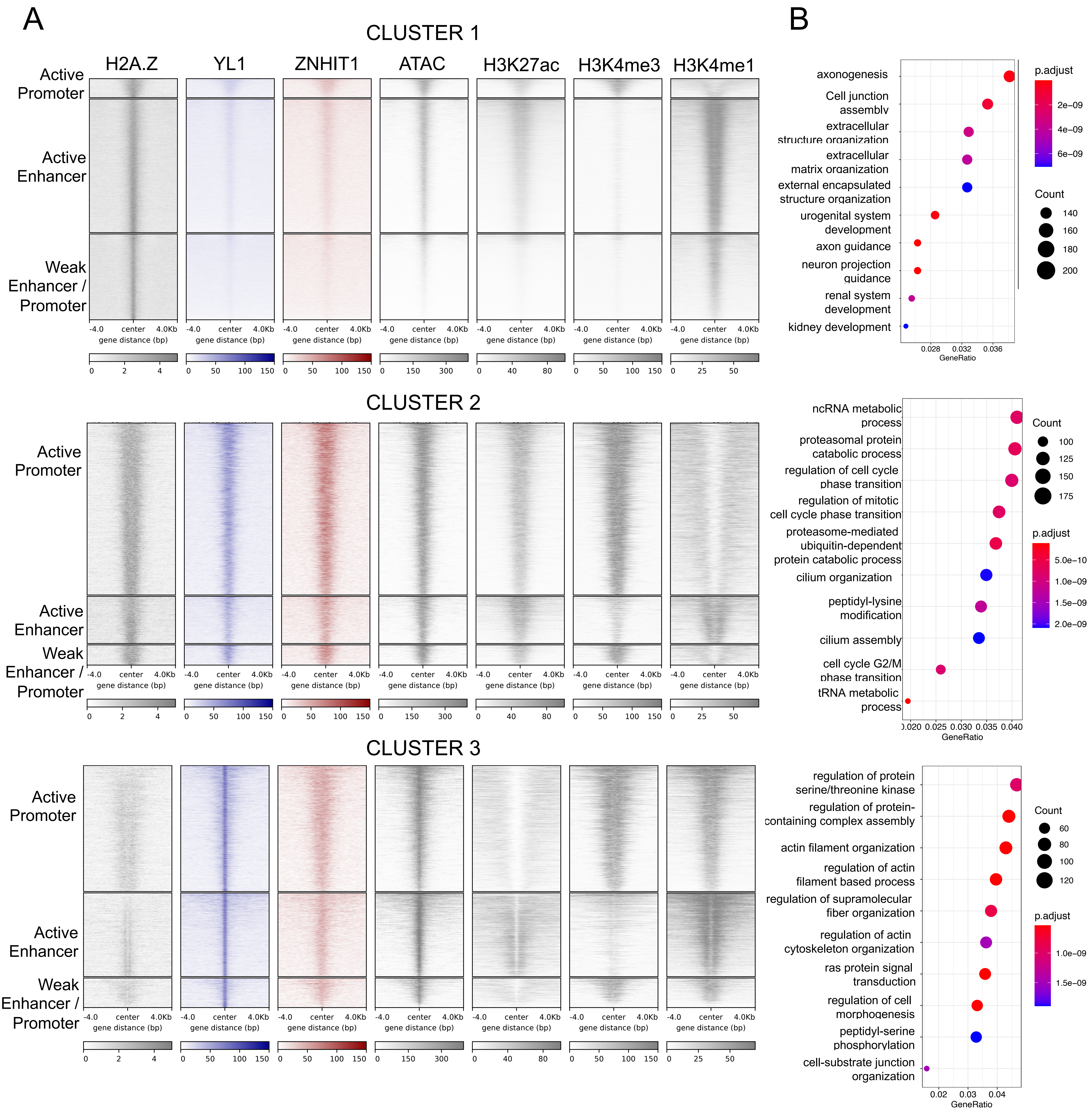

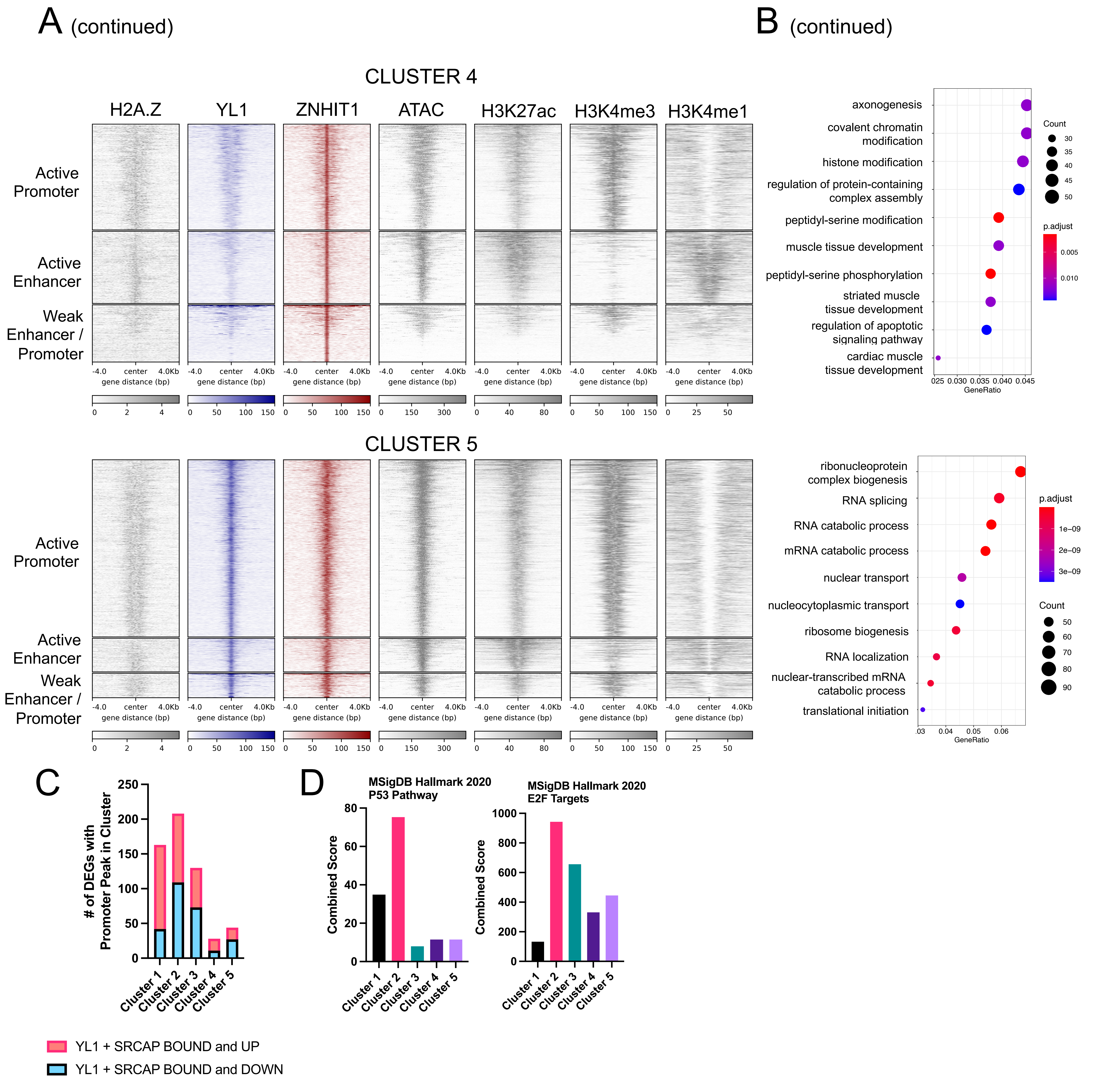


**Supp. Figure 6:** A) Heatmap of H2A.Z, YL1, ZNHIT1, ATAC-seq, H3K27ac, H3K4me3 and H3K4me1 ChIP-seq signal in SK-MEL-147 cells in Clusters 1-5, stratified by genomic annotation. B) Gene Ontology analysis of genes annotated to Clusters 1-5. C) Number of genes commonly up- and down-regulated following YL1 and SRCAP knockdown that overlap with Clusters 1-5. D) Enrichment of P53 Pathway within genes commonly upregulated following YL1 and SRCAP knockdown and annotated to Clusters 1-5 (left) and enrichment of E2F Targets within genes commonly downregulated following YL1 and SRCAP knockdown and annotated to Clusters 1-5 (right).

**Jostes et al., Supp. Fig. 7:**

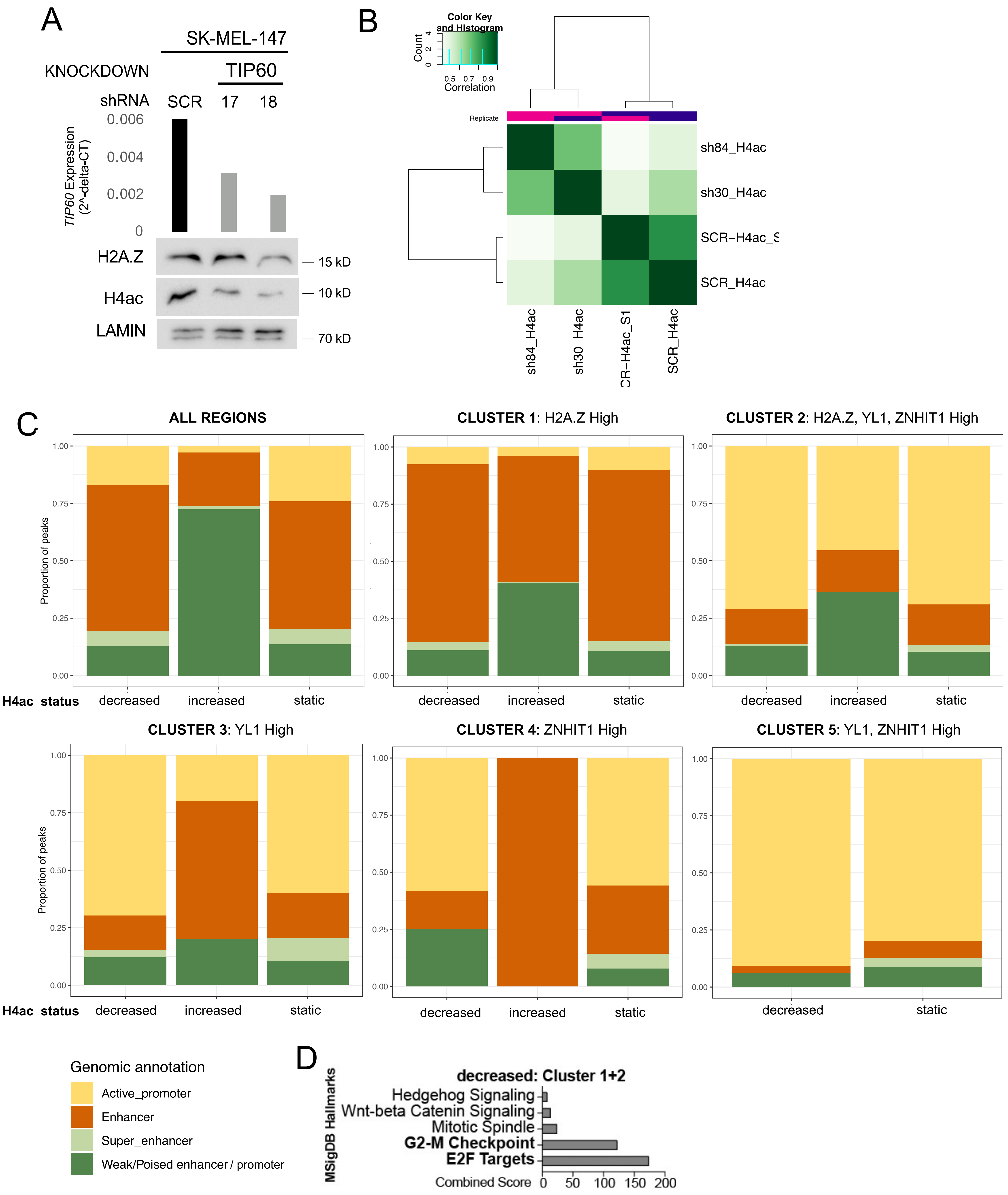


**Supp. Figure 7:** A) Top: *TIP60* mRNA expression following TIP60 knockdown compared to SCR control measured by qRT-PCR. Bottom: Corresponding Western blots showing protein levels of H2A.Z and H4ac in chromatin upon TIP60 knockdown. LAMIN serves as loading control. B) Correlation heatmap of H4ac ChIP in SCR control and YL1 knockdown samples, calculated using occupancy (peak caller score) data generated by Diffbind. C) Barplots showing gene annotations for H4ac peaks decreased, increased and static in YL1 knockdown samples versus SCR controls, stratified by Clusters as defined in Figure 3A. D) Enrichment analysis for genes with a promoter peak in Cluster 1+2 that display decreased H4 acetylation.

**Jostes et al., Supp. Fig. 8:**

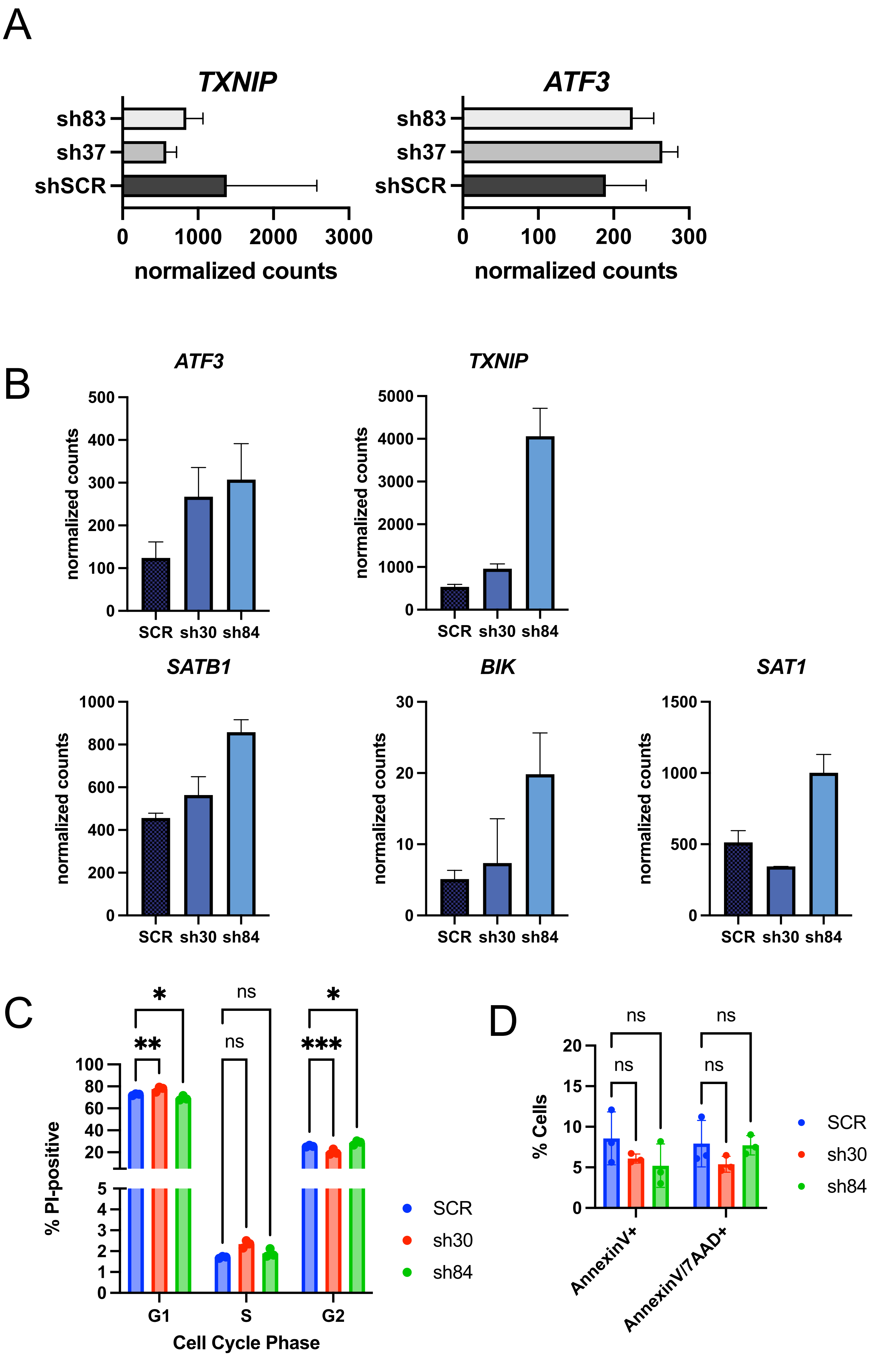


**Supp. Figure 8:** A) mRNA expression of apoptosis genes in cells infected with H2A.Z.1 (sh83) and H2A.Z.2 (sh37) shRNAs and SCR controls 6 days post-infection as measured by RNA-seq. B) mRNA expression of apoptosis genes in cells infected with YL1 shRNAs and SCR controls 3 days post-infection as measured by RNA-seq. C) Cell cycle FACS analysis of melanocytes after YL1 knockdown. Error bars indicate SD of the mean. Significance calculated using 2-way ANOVA. C) Annexin V FACS analysis of melanocytes after YL1 knockdown. Error bars indicate SD of the mean.
